# Candidate Gene Networks for Acylsugar Metabolism and Plant Defense in Wild Tomato *Solanum pennellii*

**DOI:** 10.1101/294306

**Authors:** Sabyasachi Mandal, Wangming Ji, Thomas D. McKnight

## Abstract

Many plants in the Solanaceae family secrete acylsugars, which are branched-chain and straight-chain fatty acids esterified to glucose or sucrose. These compounds have important roles in plant defense and potential commercial applications. However, several acylsugar metabolic genes remain unidentified, and little is known about regulation of this pathway. We used comparative transcriptomic analysis between low- and high-acylsugar-producing accessions of *Solanum pennellii* and found that expression levels of most acylsugar metabolic genes, including known acylsucrose biosynthetic genes and novel candidate genes (putatively encoding a ketoacyl-ACP synthase IV/II-like enzyme, peroxisomal acyl-activating enzymes, ABC transporters, and central carbon metabolic enzymes), were positively correlated with acylsugar accumulation, except two acylglucose biosynthetic genes. Genes putatively encoding oxylipin metabolic proteins, subtilisin-like proteases, and other antimicrobial defense proteins were upregulated in low-acylsugar-producing accessions, possibly to compensate for diminished defense activities of acylsugars. Gene co-expression network analysis clustered most differentially expressed genes into two separate modules and identified genetic networks associated with acylsugar production and plant defense. Transcriptome analysis after inhibition of biosynthesis of branched-chain amino acids (precursors to branched-chain fatty acids) further supported the coordinated regulation of most acylsugar candidate genes and identified three putative AP2-family transcription factor genes that form a strong co-expression network with many acylsugar metabolic genes.

## INTRODUCTION

Plants synthesize a wide variety of specialized metabolites that provide selective advantage in specific environmental conditions. Different classes of specialized metabolites are found in specific taxonomic groups and many of them are valuable phytochemicals. Acylsugars are nonvolatile and viscous specialized metabolites secreted through trichomes of solanaceous plants (Fobes et al., 1985; Kroumova et al., 2016; Moghe et al., 2017), and their roles in plant defense have been studied extensively. For example, acylsugars from *Solanum pennellii* act as feeding and oviposition deterrents for insect pests and exert toxic effects on different herbivores (Hawthorne et al., 1992; Juvik et al., 1994; Liedl et al., 1995). Acylsugars from other genera also have important roles in providing protection against herbivores and plant pathogens (Chortyk et al., 1997; Hare, 2005; Luu et al., 2017). Additionally, amphipathic acylsugar molecules are believed to reduce the surface tension of adsorbed dew, thereby providing an additional source of water for the plants (Fobes et al., 1985). Acylsugars also have potential applications as pesticides (Puterka et al., 2003), food additives (Hill and Rhode, 1999), cosmetic and personal care products (Hill and Rhode, 1999), antibiotics (Chortyk et al., 1993), and anti-inflammatory compounds (Perez-Castorena et al., 2010). Due to these important biological roles and commercial applications of acylsugars, a detailed understanding of genetic network involved in acylsugar metabolism is required for successful crop breeding programs and metabolic engineering of acylsugar production.

Acylsugars typically consist of aliphatic, branched- and/or straight-chain fatty acids esterified to hydroxyl groups of glucose or sucrose molecules. In S. *pennellii*, acylsugars are mostly 2,3,4-tri-*O*-acylated glucose esters, and some sucrose esters, with C4-C12 fatty acids (Walters and Steffens, 1990; Shapiro et al., 1994; Schilmiller et al., 2015). Predominant branched-chain fatty acids (BCFAs) include 2-methylpropanoic acid, 3-methylbutanoic acid, 2-methylbutanoic acid, and 8-methylnonanoic acid, whereas predominant straight-chain fatty acids (SCFAs) include *n*-decanoic acid and *n*-dodecanoic acid. Feeding studies demonstrated that BCFAs are derived from branched-chain amino acids (BCAAs), whereas SCFAs are hypothesized to be derived by fatty acid synthase (FAS)-mediated de novo fatty acid biosynthetic process (Walters and Steffens, 1990).

Biosynthesis of acylsugars can be broadly divided into two phases: 1) synthesis of fatty acyl chains, and 2) esterification of these acyl molecules to glucose or sucrose (Figure 1A). During acylglucose biosynthesis in S. *pennellii*, free fatty acids are initially esterified to glucose by UDP-glucose:fatty acid glucosyltransferase (UDP-Glc:FA GT) to create 1-*O*-acylglucose molecules (Ghangas and Steffens, 1993; Kuai et al., 1997). In the next step, which is catalyzed by a serine carboxypeptidase-like glucose acyltransferase (SCPL GAT), two molecules of 1-*O*-acylglucose participate in a disproportionation reaction to generate one molecule of 1,2-di-*O*-acylglucose and one molecule of glucose (Li et al., 1999; Li and Steffens, 2000). Synthesis of the final product, 2,3,4-tri-*O*-acylglucose, from 1,2-di-*O*-acylglucose requires removal of acyl chains from C1 position, and esterification or transfer of acyl chains to C3 and C4 positions of glucose. Since no tri- or tetra-acylglucose molecules are produced by the SCPL GAT, it is likely that enzymes belonging to hydrolase and/or transferase families are involved in the final steps of 2,3,4-tri-*O*-acylglucose biosynthesis (Figure 1A). In acylsucrose metabolism, acylsugar hydrolases (ASHs) and acylsugar acyltransferases (ASATs) catalyze the removal and transfer of acyl chains, respectively, (Schilmiller et al., 2012; Schilmiller et al., 2015; Fan et al., 2016; Schilmiller et al., 2016).

**Figure 1.**
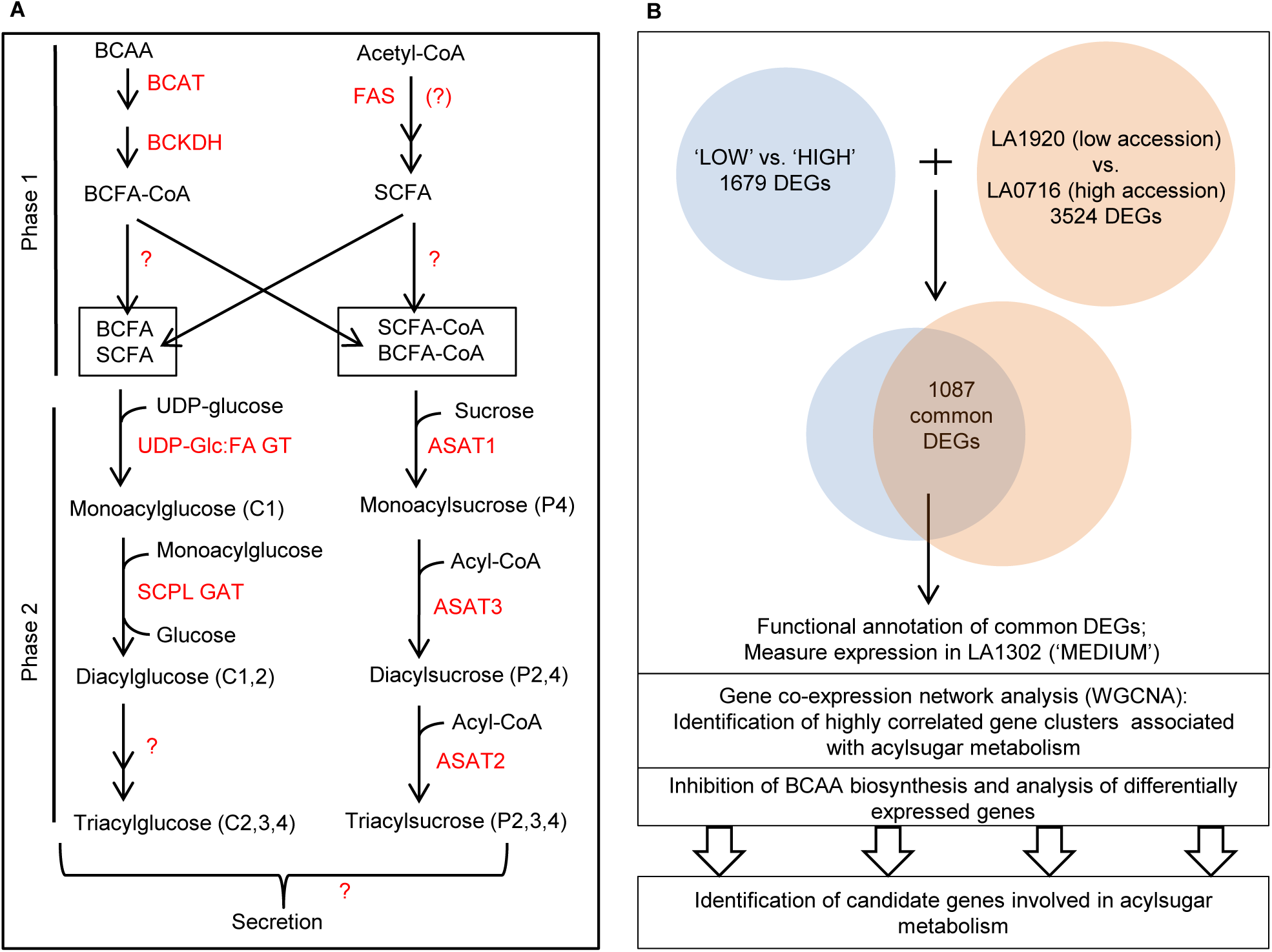
Current model of acylsugar production in *Solanum pennellii* and workflow to identify candidate genes for acylsugar metabolism. **(A)** Branched-chain fatty acids (BCFAs) are derived from branched-chain amino acids (BCAAs), and straight-chain fatty acids (SCFAs) are assumed to be produced by fatty acid synthase (FAS). UDP-glucose:fatty acid glucosyltransferase (UDP-Glc:FA GT) uses free fatty acids to initiate acylglucose biosynthesis, whereas acylsugar acyltransferase 1 (ASAT1) uses fatty acyl-CoAs to initiate acylsucrose biosynthesis. Enzymes are highlighted in red. Unidentified enzymes and transporters are marked with ‘?’. Double arrows indicate more than one enzymatic step. C1-4 indicates esterification at respective positions on glucose. P2-4 indicates esterification at respective positions on the sucrose pyranose ring. **(B)** Workflow to identify acylsugar candidate genes. Comparative transcriptomics identified 1679 differentially expressed genes (DEGs) between three low-acylsugar-producing accessions (‘LOW’) and three high-acylsugar-producing accessions (‘HIGH’). An independent comparison with higher sequencing coverage identified 3524 DEGs between another low-acylsugar-producing accession, LA1920, and the high-acylsugar-producing accession LA0716. 1087 DEGs were common to both comparisons. Expression levels of selected candidate genes were determined in LA1302, which accumulates an intermediate level of acylsugars (‘MEDIUM’). Weighted gene correlation network analysis (WGCNA) (Langfelder and Horvath, 2008) was performed to identify co-expressed gene networks associated with acylsugar accumulation. Finally, analysis of gene expression in response to inhibition of BCAA biosynthesis narrowed the list of candidates.

Acylsugars are exuded through trichomes, predominantly by type IV trichomes in S. *pennellii* (Fobes et al., 1985; Slocombe et al., 2008), and acylsucrose biosynthetic genes are expressed in trichome tip cells (Schilmiller et al., 2012; Schilmiller et al., 2015; Fan et al., 2016). However, analysis of periclinal chimeras indicates that acylglucose biosynthesis in S. *pennellii* is not trichome specific (Kuai et al., 1997). Therefore, we used leaf samples for comparative transcriptomics studies reported here. Although this approach underrepresents actual expression differences of trichome-specific genes, it ensures that a complete snapshot of candidate gene networks could be captured.

Different accessions of S. *pennellii* produce different amounts of total acylsugars (Shapiro et al., 1994). Certain accessions of S. *pennellii*, for example LA0716, can produce acylsugars up to 20% of its leaf dry weight (Fobes et al., 1985), whereas in many low-acylsugar-producing accessions, acylsugars make up less than 1% of leaf dry weight (Shapiro et al., 1994). Here, using comparative transcriptomics and gene co-expression network analysis, we exploited this natural genetic variation among *S. pennellii* accessions to identify candidate biosynthetic and regulatory genes for this trait. Comparative transcriptomics following inhibition of BCAA biosynthesis further refined the list of candidate genes and shed light on possible regulatory mechanisms of acylsugar production. Comparative transcriptomics and co-expression studies also revealed active genetic networks involved in two distinct defense pathways.

## RESULTS

### Comparative transcriptomics between low- and high-acylsugar-producing accessions

We performed differential gene expression analysis between a group of three low-acylsugar-producing accessions (LA1911, LA1912, and LA1926; collectively referred to as ‘LOW’ in this paper) and a group of three high-acylsugar-producing accessions (LA1941, LA1946, and LA0716; collectively referred to as ‘HIGH’ in this paper) of *S. pennellii* (Supplemental Table 1) (Shapiro et al., 1994). Three individual plants for each of the six accessions, with an average of >25 million processed paired-end reads for each plant, were used for this analysis (Supplemental Table 2). 19,379 genes passed our filtering criteria for minimum expression levels, and we obtained 1679 differentially expressed genes (DEGs; see Methods) (Supplemental Dataset 1: Sheet 1). Of the 1679 DEGs, 931 were upregulated and 748 were downregulated in the ‘HIGH’ group.

**Table 1.** Differentially expressed genes involved in phase 1 of acylsugar production (acyl chain synthesis).

| Gene ID | ‘LOW’ vs. ‘HIGH’ |  | LA1920 vs. LA0716 |  | Annotation |
| --- | --- | --- | --- | --- | --- |
|  | Log<sub>2</sub>FC | FDR | Log<sub>2</sub>FC | FDR |  |
| Branched-chain amino acid (BCAA)/ Branched-chain fatty acid (BCFA) metabolism |  |  |  |  |  |
| Sopen11g004560 | 1.44 | 4.30E-05 | 1.63 | 2.20E-05 | Acetolactate synthase small subunit |
| Sopen07g027240 | 1.46 | 1.00E-11 | 1.38 | 2.80E-06 | Ketol-acid reductoisomerase |
| Sopen05g032060 | 1.78 | 1.80E-10 | 1.71 | 4.50E-06 | Dihydroxy-acid dehydratase |
| Sopen08g005060 | 1.01 | 3.10E-02 | 1.38 | 3.60E-07 | Isopropylmalate synthase |
| Sopen08g005140 | 4.97 | 3.30E-13 | 5.12 | 6.40E-14 | Isopropylmalate synthase |
| Sopen04g030820 | 5.28 | 3.50E-13 | 4.55 | 4.40E-09 | Branched-chain aminotransferase-2 |
| Sopen04g026270 | 3.37 | 2.80E-12 | 2.66 | 2.10E-07 | Branched-chain keto acid dehydrogenase E1 subunit |
| Sopen01g028100 | 1.99 | 1.40E-16 | 1.53 | 9.80E-08 | Branched-chain keto acid dehydrogenase E2 subunit |
| Sopen07g023250 | 4.75 | 1.40E-08 | 3.87 | 2.30E-04 | 3-Hydroxyisobutyryl-CoA hydrolase |
| Sopen05g023470 | 1.34 | 1.40E-02 | 1.08 | 1.00E-02 | 3-Hydroxyisobutyryl-CoA hydrolase |
| Sopen12g032690 | 2.02 | 7.80E-03 | 1.87 | 1.30E-02 | Mitochondrial acyl-CoA thioesterase |
| Fatty acid synthase (FAS) components; * misannotated as two separate genes |  |  |  |  |  |
| Sopen12g004230<br>* | 5.28 | 4.40E-18 | 5.19 | 1.40E-12 | Beta-ketoacyl-ACP synthase II (N-terminus) |
| Sopen12g004240<br>* | 4.73 | 1.50E-26 | 4.39 | 8.90E-14 | Reverse transcriptase; KAS IV/ KAS II-like domain (C-terminus) |
| Sopen08g002520 | 4.56 | 1.40E-13 | 3.67 | 3.70E-07 | Beta-ketoacyl-ACP synthase III |
| Sopen05g009610 | 3.32 | 3.80E-11 | 2.73 | 1.80E-08 | Beta-ketoacyl-ACP reductase |
| Sopen12g029240 | 3.34 | 1.60E-09 | 2.85 | 2.90E-08 | Enoyl-ACP reductase domain |
| Acyl-activating enzymes |  |  |  |  |  |
| Sopen02g027670 | 3.08 | 5.50E-10 | 2.49 | 2.70E-05 | Acyl-activating enzyme 1 |
| Sopen02g027680 | 5.29 | 5.90E-12 | 4.23 | 3.70E-06 | Acyl-activating enzyme 1 |
| Sopen07g023200 | 4.16 | 2.80E-07 | 3.72 | 5.30E-05 | Acyl-activating enzyme 1 |
| Sopen07g023220 | 5.81 | 3.80E-27 | 4.02 | 3.60E-06 | Acyl-activating enzyme 1 |

1 Log<sub>2</sub>FC and FDR indicate log<sub>2</sub> (fold-change) and false discovery rate (*P* values adjusted for 2 multiple-testing), respectively. Positive and negative log<sub>2</sub>FC values indicate higher and lower 3 expression levels, respectively in high-acylsugar-producing accessions.

**Table 2.**
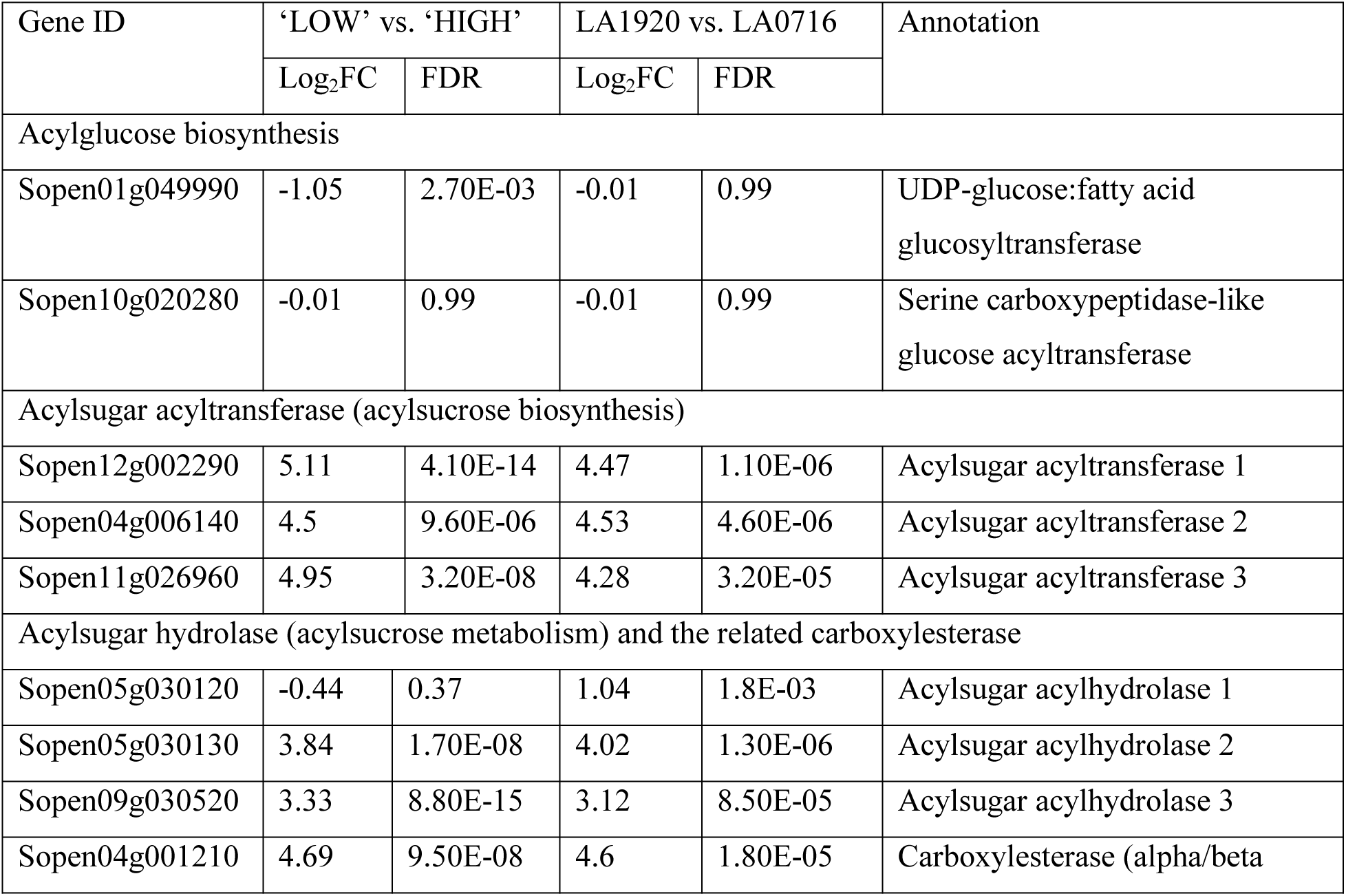
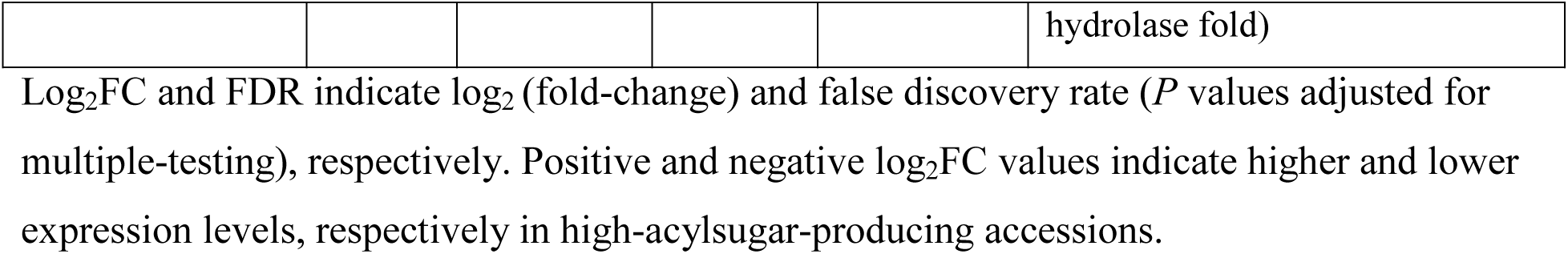
Genes involved in phase 2 of acylsugar production.

| Gene ID | ‘LOW’ vs. ‘HIGH’ |  | LA1920 vs. LA0716 |  | Annotation |
| --- | --- | --- | --- | --- | --- |
|  | Log<sub>2</sub>FC | FDR | Log<sub>2</sub>FC | FDR |  |
| Acylglucose biosynthesis |  |  |  |  |  |
| Sopen01g049990 | -1.05 | 2.70E-03 | -0.01 | 0.99 | UDP-glucose:fatty acid glucosyltransferase |
| Sopen10g020280 | -0.01 | 0.99 | -0.01 | 0.99 | Serine carboxypeptidase-like glucose acyltransferase |
| Acylsugar acyltransferase (acylsucrose biosynthesis) |  |  |  |  |  |
| Sopen12g002290 | 5.11 | 4.10E-14 | 4.47 | 1.10E-06 | Acylsugar acyltransferase 1 |
| Sopen04g006140 | 4.5 | 9.60E-06 | 4.53 | 4.60E-06 | Acylsugar acyltransferase 2 |
| Sopen11g026960 | 4.95 | 3.20E-08 | 4.28 | 3.20E-05 | Acylsugar acyltransferase 3 |
| Acylsugar hydrolase (acylsucrose metabolism) and the related carboxylesterase |  |  |  |  |  |
| Sopen05g030120 | -0.44 | 0.37 | 1.04 | 1.8E-03 | Acylsugar acylhydrolase 1 |
| Sopen05g030130 | 3.84 | 1.70E-08 | 4.02 | 1.30E-06 | Acylsugar acylhydrolase 2 |
| Sopen09g030520 | 3.33 | 8.80E-15 | 3.12 | 8.50E-05 | Acylsugar acylhydrolase 3 |
| Sopen04g001210 | 4.69 | 9.50E-08 | 4.6 | 1.80E-05 | Carboxylesterase (alpha/beta hydrolase fold) |

Log<sub>2</sub>FC and FDR indicate log<sub>2</sub> (fold-change) and false discovery rate (*P* values adjusted for multiple-testing), respectively. Positive and negative log<sub>2</sub>FC values indicate higher and lower expression levels, respectively in high-acylsugar-producing accessions.

We also compared transcriptomes between another low-acylsugar-producing accession, LA1920, and the high-acylsugar-producing accession LA0716, with each accession having four biological replicates and slightly higher sequencing coverage, as an independent examination of differential gene expression. In this analysis, 19,967 genes passed our filtering criteria for minimum expression levels (see Methods), and we identified 3524 DEGs (1465 upregulated and 2059 downregulated genes in LA0716) (Supplemental Dataset 1: Sheet 2). To identify candidate genes most likely to be involved in acylsugar metabolism, we compared ‘LOW’ vs. ‘HIGH’ DEGs with LA1920 vs. LA0716 DEGs (Figure 1B). 1087 DEGs were common to both comparisons (586 upregulated and 501 downregulated genes in high-acylsugar-producing accessions; Supplemental Dataset 1: Sheet 3). Analysis of Gene Ontology (GO) terms associated with the annotated DEGs revealed that genes involved in fatty acid metabolism, regulation of transcription and translation, as well as genes whose products are predicted to localize to chloroplasts (the location for BCAA and SCFA biosynthesis), were strongly represented in the 1087 DEGs (Supplemental Figure 1).

We also examined the leaf transcriptome of accession LA1302 (three biological replicates), which accumulates an intermediate level of acylsugars (referred to as ‘MEDIUM’ in this paper; Supplemental Table 1) (Shapiro et al., 1994), to determinate whether expression levels of candidate genes were consistent with the amount of acylsugar production (Figure 1B). Multi-dimensional scaling plots showed a clear distinction in gene expression profiles among groups of accessions that accumulate different amounts of acylsugars (Supplemental Figure 2A and 2B). Volcano plots showed high log_2_fold-change values associated with low false discovery rate (FDR) values, indicating robust differential expression (Supplemental Figure 2C and 2D). On the other hand, ten ‘housekeeping genes’ showed similar expression levels across all biological groups (Supplemental Figure 3).

**Figure 2.**
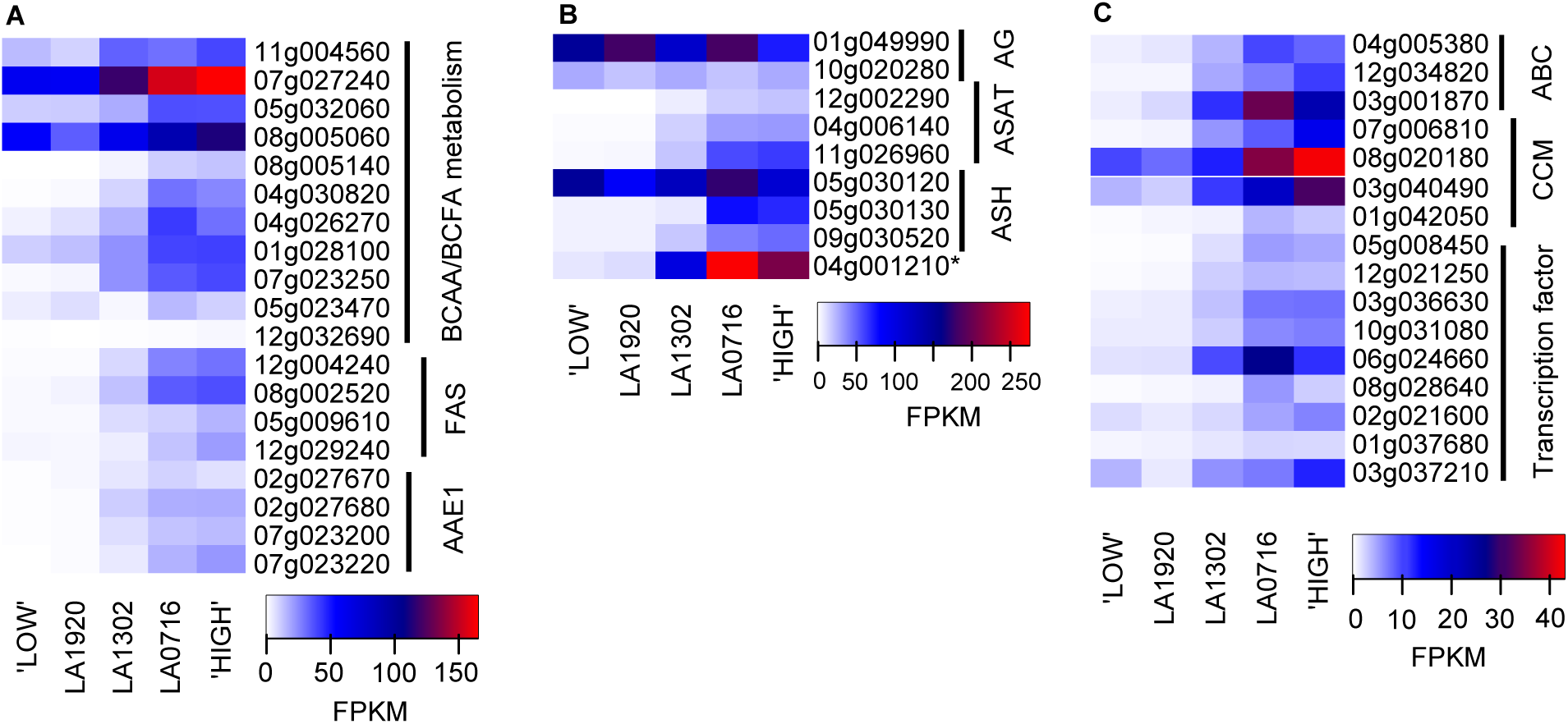
Heatmaps showing expression levels of genes with known and putative functions in acylsugar metabolism. Genes are designated by their gene identifier numbers (Sopen IDs). **(A)** Acylsugar phase 1-related genes (Table 1). Expression levels of many candidate genes in LA1302 (‘MEDIUM’ accession) were intermediate between low- and high-acylsugar-producing accessions. **(B)** Acylsugar phase 2-related genes (Table 2). *Sopen04g001210* (marked with an asterisk) was identified as a carboxylesterase gene related to ASH, but the encoded protein does not exhibit in vitro activity against tomato acylsucrose fractions (Schilmiller et al., 2016). **(C)** Other genes related to acylsugar metabolism (Table 3). Two genes *(Sopen04g023150* and *Sopen01g047950)* were removed from the heatmap due to their very high expression levels. *Sopen04g023150* had FPKM values of 570, 624, 455, 179, and 254 in ‘LOW’, LA1920, LA1302, LA0716, and ‘HIGH’ groups, respectively. *Sopen01g047950* had FPKM values of 61, 203, 79, 29, and 19 in ‘LOW’, LA1920, LA1302, LA0716, and ‘HIGH’ groups, respectively.Abbreviations: FPKM= fragments per kilobase of transcript per million mapped reads; BCAA= branched-chain amino acid; FAS= fatty acid synthase components; AAE1= acyl-activating enzyme 1; AG= acylglucose phase 2; ASAT= acylsucrose acyltransferase; ASH= acylsugar hydrolase; ABC= ABC transporters; CCM= central carbon metabolism.

**Figure 3.**
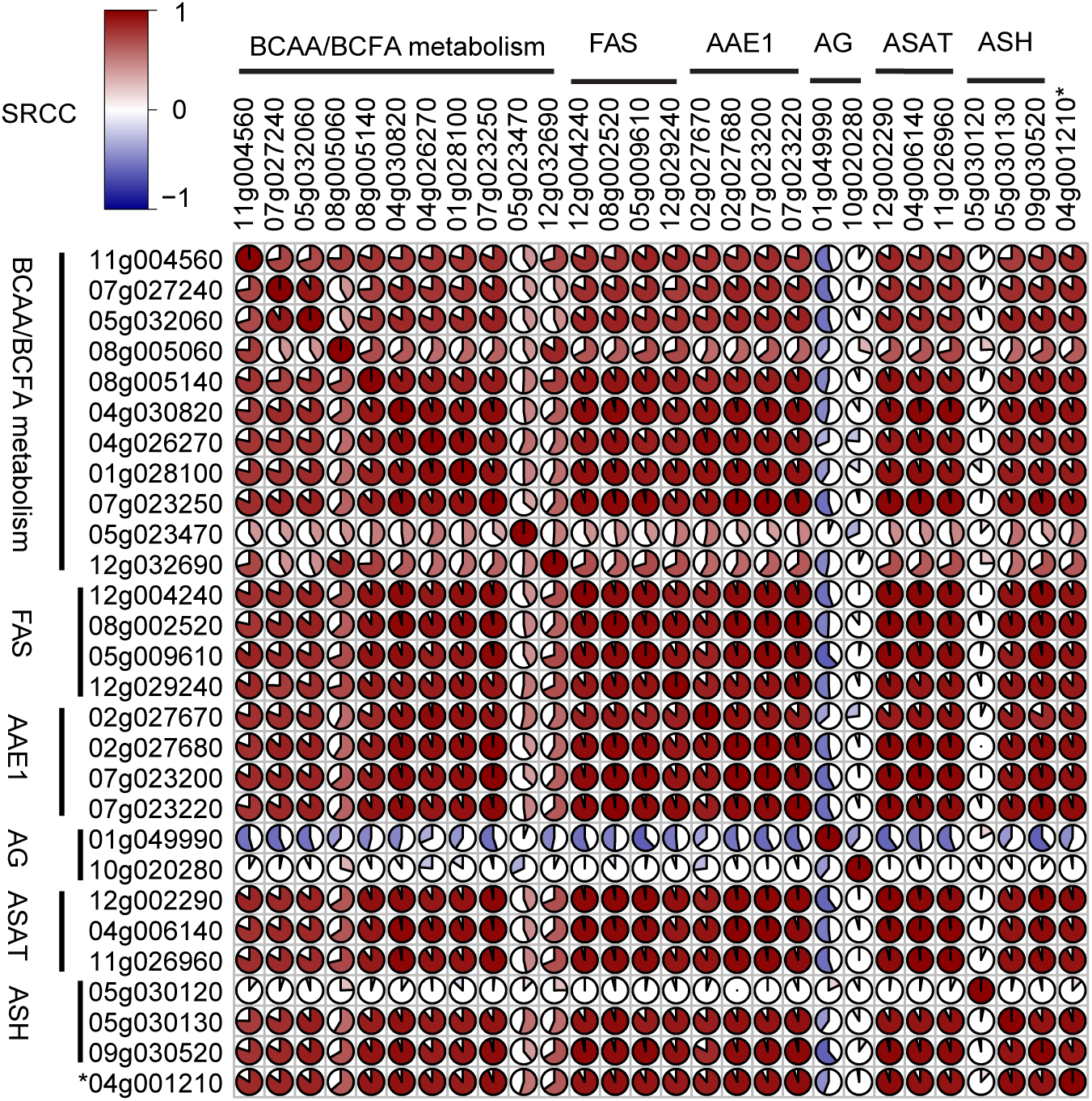
Correlation among expression profiles of selected genes. FPKM values for 28 phase 1- and phase 2-related genes (from Table 1 and Table 2, respectively) in our 29 samples were used to determine pairwise Spearman’s rank correlation coefficients (SRCC). A [28x28] matrix was created to visualize correlation. SRCC values and associated P values are given in Supplemental Dataset 2. *Sopen04g001210* (marked with an asterisk) was identified as a carboxylesterase gene related to ASH (Schilmiller et al., 2016). Abbreviations: BCAA= branched-chain amino acid; FAS= fatty acid synthase components; AAE1= acyl-activating enzyme 1; AG= acylglucose phase 2; ASAT= acylsucrose acyltransferase; ASH= acylsugar hydrolase.

### Branched-chain amino acid (BCAA) metabolic genes are upregulated in high-acylsugar-producing accessions

Because BCAAs are precursors to acylsugar BCFAs (Walters and Steffens, 1990), we examined genes involved in BCAA metabolism. Many genes involved in both biosynthesis of BCAAs in plastids and conversion of BCAAs to branched-chain acyl molecules in mitochondria (Binder, 2010; Maloney et al., 2010) were upregulated in high-acylsugar-producing accessions (Table 1; Supplemental Figure 4). Most of these BCAA/BCFA metabolic DEGs showed intermediate levels of expression in the ‘MEDIUM’ accession LA1302 (Figure 2A). Correlation analysis (Spearman’s rank correlation coefficient, SRCC; denoted by ρ) using gene expression profiles in our 29 samples revealed strong positive correlation among most of these genes (Figure 3).

**Table 3.**
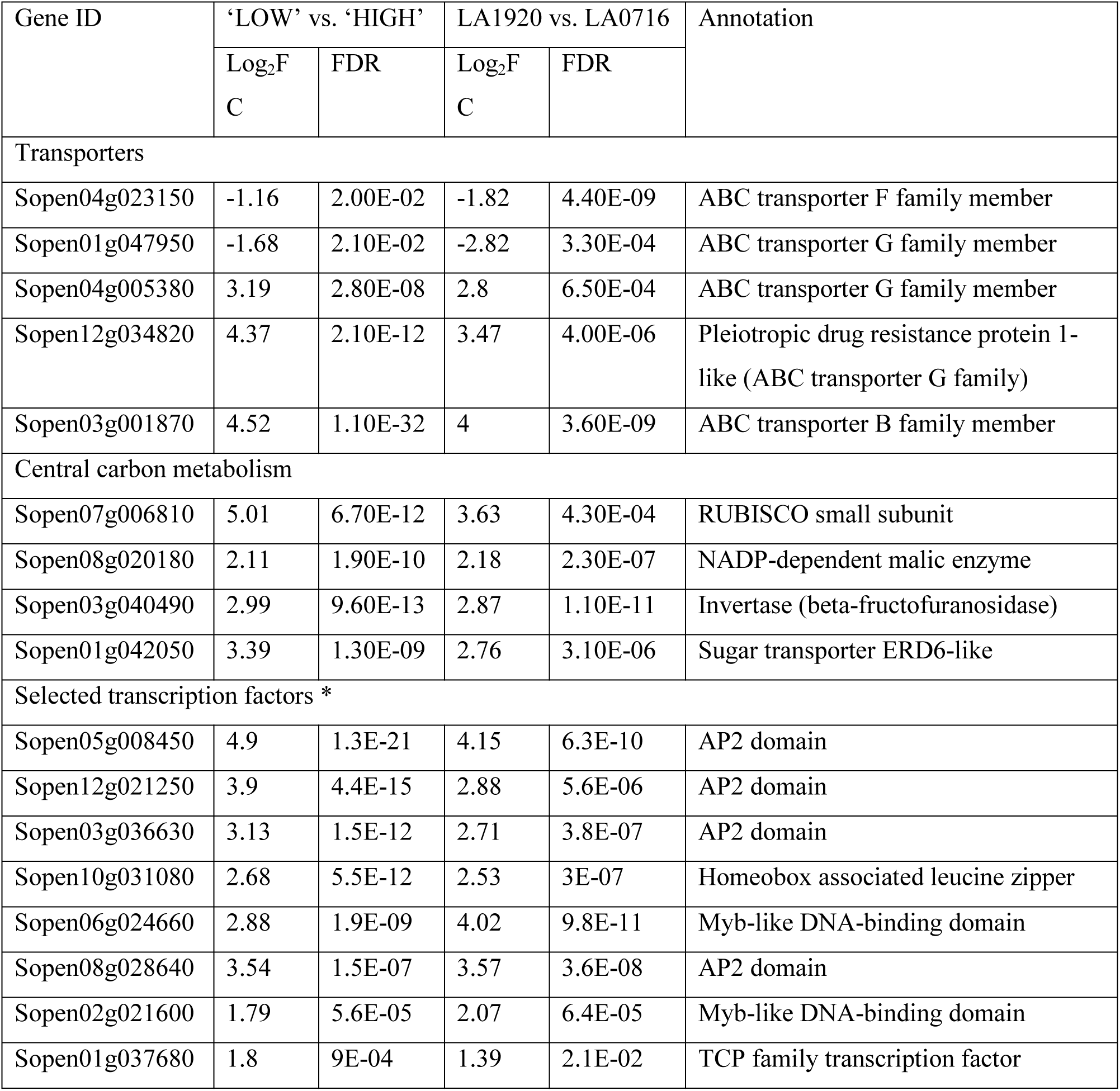
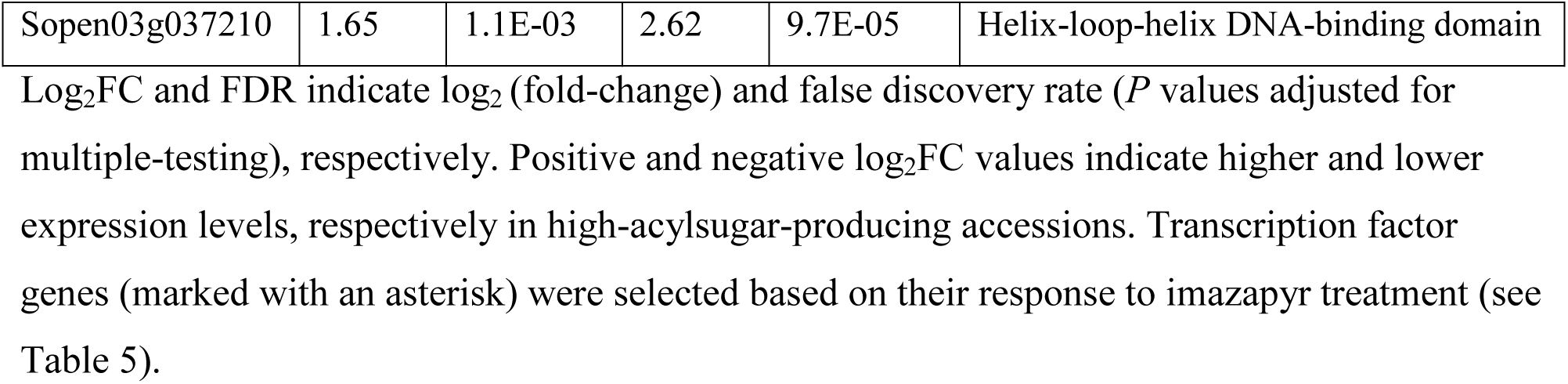
Differentially expressed genes involved in transport process, central carbon metabolism, and regulation of gene expression.

**Figure 4.**
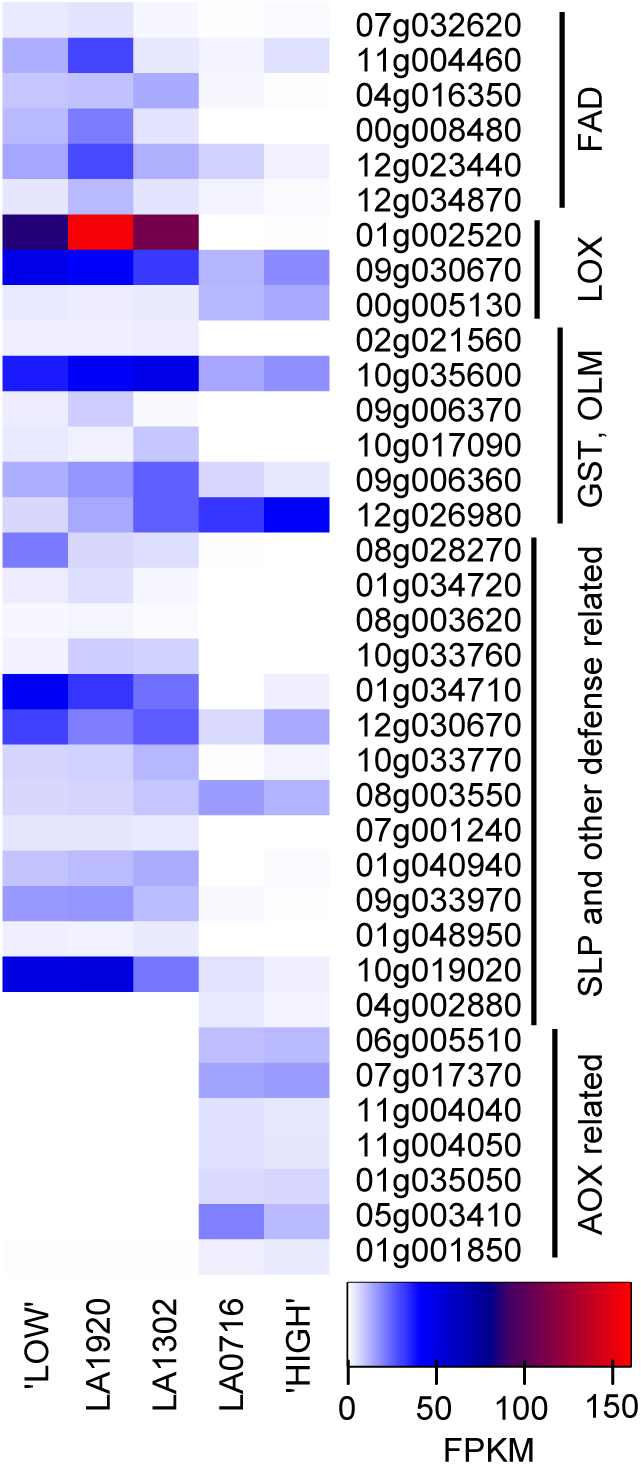
Heatmap showing expression levels of genes putatively involved in plant defense (Table 4 genes). Expression levels of many genes in LA1302 were similar to low-acylsugar-producing accessions but different from high-acylsugar-producing accessions. Some oxylipin-related defense genes, which were upregulated in low-acylsugar-producing accessions, had higher expression levels than amine oxidase-related defense genes, which were upregulated in high-acylsugar-producing accessions. Abbreviations: FPKM= fragments per kilobase of transcript per million mapped reads; FAD= fatty acid desaturase; LOX= lipoxygenase; GST= glutathione S-transferase; OLM= oxylipin metabolism; SLP= subtilisin-like protease; AOX= amine oxidase.

### Genes putatively encoding a KAS IV/ KAS II-like enzyme and other FAS components are upregulated in high-acylsugar-producing accessions

*Sopen12g004230*, predicted to encode a beta-ketoacyl-(acyl-carrier-protein) synthase II (KAS II)-like enzyme, had 39-fold higher expression in the ‘HIGH’ group (FDR= 4.40E-18). The SCFA profile of *S. pennellii* acylsugars (predominantly C10 and C12, with a trace amount of C8 in LA1302) is strikingly similar to that of the seed oil in many *Cuphea* species (Shapiro et al., 1994; Dehesh et al., 1998; Slabaugh et al., 1998; Schutt et al., 2002). A specialized version of KAS II, referred to as KAS IV, is an important determinant of chain length in *Cuphea* medium-chain (C8-C12) fatty acids (Dehesh et al., 1998; Slabaugh et al., 1998; Schutt et al., 2002). We compared amino acid sequences of five KAS IV/ KAS II-like enzymes from four *Cuphea* species with the Sopen12g004230 sequence, but no similarity was found. However, sequence analysis of the adjacent Sopen12g004240 coding region (1785-aa; annotated as reverse transcriptase) revealed a KAS domain at the C-terminal end (1374-1784), and BLAST analysis of *Cuphea* KAS IV/ KAS II-like enzymes showed high sequence similarity with Sopen12g004240 (77%-83% identity, 87%-92% similarity, 82%-90% coverage, e-value 0.0; Supplemental Table 3). Apparently, a transposon inserted into an intron of *Sopen12g004230-Sopen12g004240*, which led to misannotation of this locus as two separate genes. We confirmed that this locus produces a single transcript with RT-PCR analysis (Supplemental Figure 5). Blauth et al. (1999) used an intraspecific F_2_ population derived from the cross between accessions LA0716 and LA1912, to identify a quantitative trait locus on the upper arm of chromosome 12 that is associated with C10 SCFA production (Blauth et al., 1999). The locus containing *Sopen12g004230-Sopen12g004240* is in this region.

**Table 4.** Differentially expressed genes putatively involved in oxylipin-related and amine oxidase-related defense systems.

| Gene ID | ‘LOW’ vs. ‘HIGH’ |  | LA1920 vs. LA0716 |  | Annotation |
| --- | --- | --- | --- | --- | --- |
|  | Log<sub>2</sub>FC | FDR | Log<sub>2</sub>FC | FDR |  |
| Desaturase and lipoxygenase |  |  |  |  |  |
| Sopen07g032620 | -2.43 | 2.10E-08 | -3.12 | 4.10E-09 | Fatty acid desaturase 4 |
| Sopen11g004460 | -1.37 | 2.30E-03 | -3.86 | 1.40E-30 | Stearoyl-(ACP)-9-desaturase |
| Sopen04g016350 | -4.23 | 7.00E-22 | -3.00 | 2.30E-05 | Delta(12)-fatty acid desaturase |
| Sopen00g008480 | -6.29 | 9.10E-08 | -8.72 | 4.60E-67 | Delta(12)-fatty acid desaturase |
| Sopen12g023440 | -2.71 | 5.60E-05 | -2.03 | 1.60E-02 | Delta(12)-fatty acid desaturase |
| Sopen12g034870 | -2.69 | 4.10E-04 | -2.48 | 1.00E-02 | Delta(12)-fatty acid desaturase |
| Sopen01g002520 | -7.99 | 2.40E-15 | -10.7 | 2.30E-40 | Linoleate 13S-lipoxygenase |
| Sopen09g030670 | -1.38 | 3.30E-08 | -1.84 | 1.90E-10 | Linoleate 9S-lipoxygenase |
| Sopen00g005130 | 2.03 | 9.70E-09 | 1.97 | 1.00E-05 | Linoleate 9S-lipoxygenase |
| Oxylipin signaling and detoxification |  |  |  |  |  |
| Sopen02g021560 | -3.55 | 6.80E-10 | -4.35 | 3.40E-10 | Transcription factor TGA5-like |
| Sopen10g035600 | -1.06 | 5.10E-05 | -1.63 | 5.50E-11 | 12-Oxo-phytodienoate reductase |
| Sopen09g006370 | -7.11 | 2.20E-16 | -11.1 | 1.40E-08 | Glutathione S-transferase-like |
| Sopen10g017090 | -4.89 | 7.00E-09 | -7.08 | 7.10E-15 | Glutathione S-transferase |
| Sopen09g006360 | -1.83 | 2.10E-03 | -1.42 | 3.30E-02 | Glutathione S-transferase |
| Sopen12g026980* | 2.72 | 1.80E-09 | 1.21 | 3.40E-04 | Glutathione S-transferase |
| Other plant defense-related |  |  |  |  |  |
| Sopen08g028270 | -7.03 | 2.90E-16 | -4.15 | 8.60E-05 | Subtilisin-like protease precursor |
| Sopen01g034720 | -8.23 | 5.90E-15 | -10.5 | 5.70E-21 | Subtilisin-like serine protease SBT4 |
| Sopen08g003620 | -3.86 | 6.40E-12 | -4.11 | 7.10E-07 | Subtilisin-like protease SBT1.7 |
| Sopen10g033760 | -2.98 | 9.70E-05 | -8.04 | 3.10E-08 | Subtilisin-like protease SBT1.7 |
| Sopen01g034710 | -4.13 | 6.80E-03 | -9.72 | 4.50E-31 | Subtilisin-like protease SBT1.7 |
| Sopen12g030670 | -1.13 | 7.60E-03 | -1.76 | 2.00E-06 | Subtilisin-like protease SBT1.7 |
| Sopen10g033770 | -1.77 | 1.80E-02 | -3.70 | 4.90E-09 | Subtilisin-like protease SBT1.7 |
| Sopen08g003550 | 1.02 | 1.70E-07 | 1.18 | 1.80E-04 | Subtilisin-like protease SBT1.7 |
| Sopen07g001240 | -5.34 | 1.00E-09 | -5.30 | 8.90E-07 | Chitotriosidase-1 (chitinase activity) |
| Sopen01g040940 | -3.41 | 2.80E-04 | -6.03 | 1.20E-08 | Wound-induced protein WIN1-like (chitin binding) |
| Sopen09g033970 | -5.43 | 1.70E-14 | -3.95 | 3.60E-06 | Pathogenesis-related protein STH-2 |
| Sopen01g048950 | -3.72 | 4.20E-05 | -4.92 | 2.30E-05 | Pathogenesis-related protein 1A-like |
| Sopen10g019020 | -4.16 | 3.80E-07 | -3.42 | 4.80E-12 | Kirola-like defense response protein |
| Sopen04g002880 | 6.04 | 1.70E-15 | 4.14 | 9.50E-05 | Kirola-like defense response protein |
| Amine oxidase-related defense system |  |  |  |  |  |
| Sopen06g005510 | 7.52 | 2.60E-42 | 6.46 | 1.90E-17 | Glutamine synthetase catalytic domain |
| Sopen07g017370 | 7.54 | 3.20E-47 | 6.86 | 3.90E-16 | Copper-containing amine oxidase |
| Sopen11g004040 | 7.30 | 9.20E-15 | 6.89 | 3.20E-18 | Protein FLOWERING LOCUS D (flavin-containing amine oxidase) |
| Sopen11g004050 | 6.04 | 2.30E-14 | 4.53 | 1.20E-15 | Protein FLOWERING LOCUS D (flavin-containing amine oxidase) |
| Sopen01g035050 | 7.84 | 5.00E-57 | 6.07 | 6.60E-13 | Aldehyde oxidase |
| Sopen05g003410 | 6.95 | 2.00E-22 | 6.11 | 6.00E-09 | Extensin-like protein |
| Sopen01g001850 | 2.97 | 2.10E-10 | 2.67 | 1.00E-06 | Extensin-like protein |

**Figure 5.**
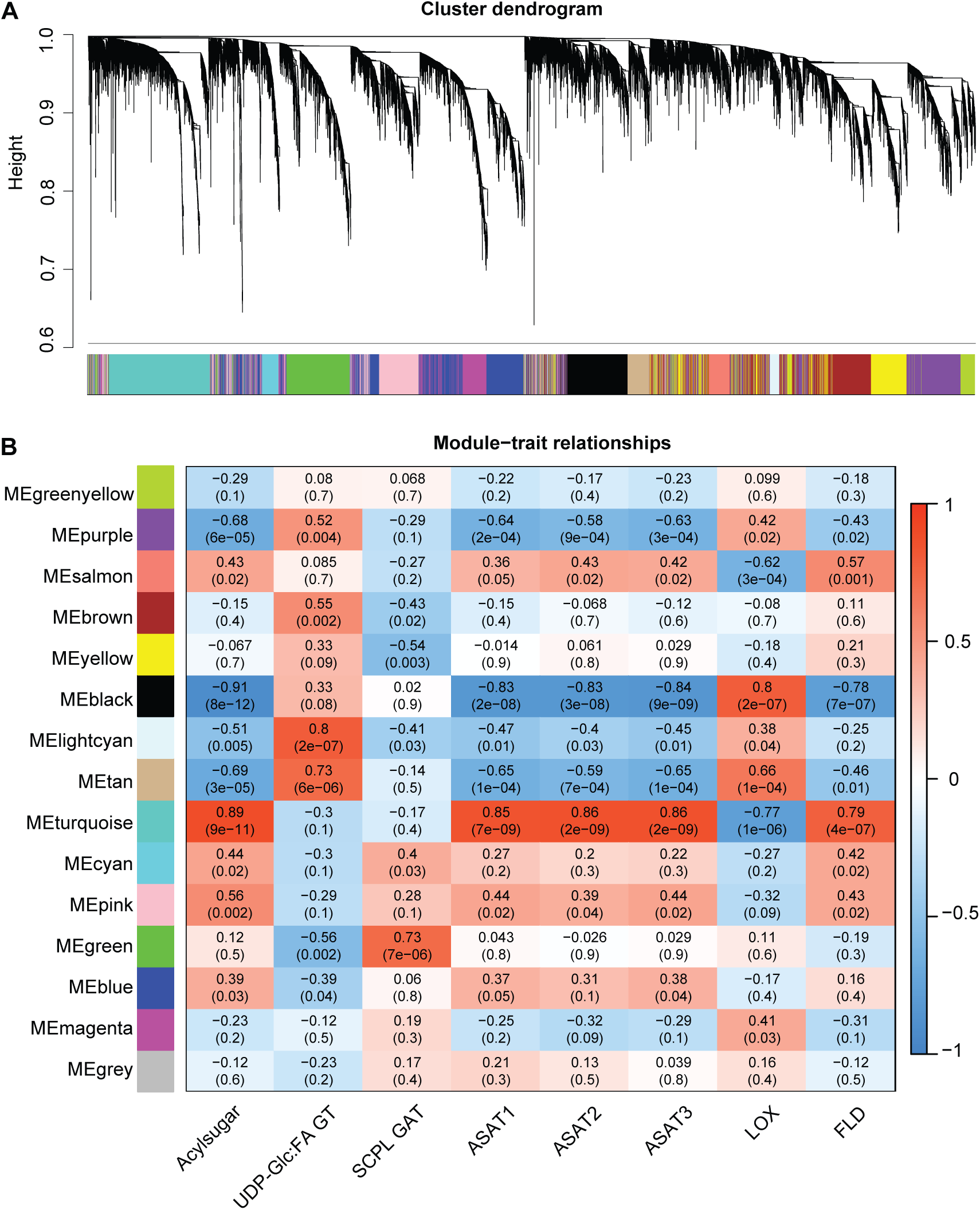
Weighted gene correlation network analysis (WGCNA) (Langfelder and Horvath, 2008). **(A)** Dendogram clustering 19,378 genes into 15 co-expressed modules based on their expression profiles in all 29 biological samples (see Methods section). For each module, a module eigengene was selected by WGCNA as representative of the expression profile for that module. **(B)** Association of module eigengenes (MEs) with acylsugar level and seven selected gene expression profiles. The seven genes encode acylglucose biosynthetic enzymes (UDP-Glc:FA GT and SCPL GAT), acylsucrose biosynthetic enzymes (ASAT1, ASAT2, and ASAT3), and defense related proteins (LOX and FLD as representatives of oxylipin-related and amine oxidase-related defense networks, respectively). Spearman’s rank correlation coefficient (SRCC) values between MEs and traits are shown; P values associated with SRCC are given in parentheses.Abbreviations: UDP-Glc:FA GT= UDP-glucose:fatty acid glucosyltransferase; SCPL GAT= serine carboxypeptidase-like glucose acyltransferase; ASAT= acylsugar acyltransferase; LOX= lipoxygenase; FLD= FLOWERING LOCUS D.

DEGs putatively encoding other components of the FAS complex were also upregulated in high-acylsugar-producing accessions (Table 1). Expression patterns of these genes and *Sopen12g004240* (KAS IV/ KAS II-like) showed very strong positive correlation among themselves, as well as with BCAA metabolic DEGs (Figure 3; Supplemental Dataset 2).

### Expression patterns of four genes putatively encoding acyl-activating enzymes show very strong positive correlation with FAS components

Acyl-activating enzymes (AAEs) are ATP/AMP-binding proteins that activate different carboxylic acids by forming fatty acyl-CoA molecules from free fatty acids, ATP and CoA (Shockey et al., 2003). We found four DEGs putatively encoding acyl-activating enzyme 1 (AAE1), and they had 8-to 56-fold higher expression in the ‘HIGH’ group (FDR= 2.80E-07 to 3.80E-27; Table 1). In *Arabidopsis thaliana*, AAE7 and AAE11, which belong to the same phylogenetic clade as AAE1, show selective activity against short- and medium-chain fatty acids (Shockey et al., 2003); interestingly, these four AAE1 genes exhibited very strong positive correlation with FAS components (*ρ*= 0.87 to 0.98; *P* = 1.07E-9 to 4.44E-16; Figure 3).

### Acylsucrose metabolic genes are upregulated in high-acylsugar-producing accessions and show strong positive correlation with phase 1 DEGs

Three genes encoding acylsugar acyltransferases (ASATs), which are involved in phase 2 of acylsucrose biosynthesis (Schilmiller et al., 2015; Fan et al., 2016; Fan et al., 2017), were upregulated in high-acylsugar-producing accessions (Table 2). ASATs use acyl-CoA molecules as their substrates (Schilmiller et al., 2012; Schilmiller et al., 2015; Fan et al., 2016; Fan et al., 2017), and free straight-chain fatty acids (SCFA) produced by FAS-catalyzed de novo fatty acid biosynthesis must be activated to their acyl-CoA derivatives (SCFA-CoA) before they can be used by ASATs (Figure 1A). Interestingly, all three ASAT genes exhibited very strong positive correlation with both AAE1 genes and FAS components (Figure 3; Supplemental Dataset 2). Using data of Ning et al. (2015), who published expression profiles of *S. lycopersicum* genes in isolated stem trichomes versus shaved stems, we also investigated if AAE1 homologs in S. *lycopersicum* show trichome-enriched expression pattern, since ASAT genes are expressed in apical cells of type I/IV glandular trichomes (Schilmiller et al., 2012; Schilmiller et al., 2015; Fan et al., 2016). We identified *Solyc02g082870, Solyc02g082880, Solyc07g043630*, and *Solyc07g043660* as putative orthologs of the four AAE1 DEGs *Sopen02g027670, Sopen02g027680, Sopen07g023200*, and *Sopen07g023220*, respectively (Supplemental Dataset 3). According to Ning et al. (2015), these four tomato genes show 300-, 26-, 562-, and 294-fold higher expression levels, respectively, in isolated stem trichomes compared to shaved stems. In fact, three of these AAE1 homologs show almost no expression in shaved stems, but high expression levels in isolated trichomes, similar to ASAT genes.

Recently, three genes encoding acylsugar hydrolases (ASHs; carboxylesterases that remove acyl groups from acylsucrose molecules) and a related carboxylesterase gene *(Sopen04g001210)* were reported in cultivated and wild tomato (Schilmiller et al., 2016). These genes, except *Sopen05g030120* (Sp-ASH1), were in the list of 1087 common DEGs, and showed strong positive correlation with genes putatively involved in phase 1 acyl chain synthesis (Figure 3; Supplemental Dataset 2). The expression profile of *ASH1* in different tissues of S. *lycopersicum* led the authors to propose additional non-trichome-localized functions for Sl-ASH1 (Schilmiller et al., 2016); our results further support the idea that ASH1 may not have a critical role in trichome acylsucrose metabolism.

### Expression levels of two acylglucose biosynthetic genes are not consistent with acylsugar levels

The first two steps in phase 2 of acylglucose biosynthesis are catalyzed by UDP-Glc:FA GT and SCPL GAT, respectively (Figure 1A) (Ghangas and Steffens, 1993; Kuai et al., 1997; Li et al., 1999; Li and Steffens, 2000). Interestingly, *Sopen01g049990*, encoding the UDP-Glc:FA GT, showed 2-fold higher expression in the ‘LOW’ group (FDR= 2.70E-03), whereas *Sopen10g020280*, which encodes the SCPL GAT, had no significant difference in expression between low- and high-acylsugar-producing accessions (Table 2). Similarly, *Sopen01g049990* showed weak negative correlation with genes putatively involved in acyl chain synthesis (phase 1), whereas *Sopen10g020280* showed no significant correlation (Figure 3; Supplemental Dataset 2).

The UDP-Glc:FA GT uses free fatty acids (such as 2-methypropanoate), but not their CoA derivatives, to initiate acylglucose biosynthesis (Ghangas and Steffens, 1993). In mitochondria, activity of the branched-chain keto acid dehydrogenase complex forms BCFA-CoA derivatives of BCAAs, and these acyl-CoA molecules therefore must be released as free fatty acids (BCFAs) by thioesterases before they could be used for acylglucose biosynthesis (Figure 1A). One gene *(Sopen12g032690)*, predicted to encode a mitochondrial acyl-CoA thioesterase, had 4-fold higher expression in the ‘HIGH’ group (FDR= 7.80E-03; Table 1).

### Three genes putatively encoding ATP-binding cassette (ABC) transporters are upregulated in high-acylsugar-producing accessions

One important question in acylsugar metabolism is how acylsugar molecules are exported out of trichome cells. It remains unclear whether acylsugars are transported across the plasma membrane via membrane-associated transporter proteins or via vesicular transport. Five genes belonging to the ATP-binding cassette (ABC) transporter family showed differential expression, and three of them *(Sopen12g034820, Sopen04g005380*, and *Sopen03g001870)* were upregulated in high-acylsugar-producing accessions (Table 3). We identified *Solyc12g100190, Solyc04g010200*, and *Solyc03g005860* as putative orthologs of these three ABC transporters (Supplemental Dataset 3). Based on data of Ning et al. (2015), these three tomato genes show 143-, 160-, and 470-fold, respectively higher expression in isolated trichomes than in underlying tissues, with expression of *Solyc03g005860* being very similar to that of trichome-specific ASAT genes. Because acylsugars are secreted by trichome tip cells (Fobes et al., 1985), it is reasonable to expect that genes involved in acylsugar secretion will exhibit trichome-specific/ trichome-enriched expression pattern. On the other hand, putative *S. lycopersicum* orthologs *(Solyc04g051800* and *Solyc01g105450*; Supplemental Dataset 3) of the other ABC transporter DEGs (*Sopen04g023150* and *Sopen01g047950*), which were downregulated in high-acylsugar-producing accessions, have similar expression levels in both trichomes and underlying tissues (Ning et al., 2015).

### Genes involved in central carbon metabolism are upregulated in high-acylsugar-producing accessions

In high-acylsugar-producing accessions, acylsugar accumulation represents an astounding metabolic investment (up to 20% of leaf dry weight in LA0716) (Fobes et al., 1985; Shapiro et al., 1994), and we investigated whether genes involved in central carbon metabolism were differentially expressed to support acylsugar production. Two genes *(Sopen07g006810* and *Sopen08g020180)* putatively encoding the RUBISCO small subunit and chloroplast NADP-malic enzyme, respectively were upregulated in high-acylsugar-producing accessions (Table 3). Compared to other RUBISCO small subunit genes, which were not differentially expressed and had very high expression levels (FPKM as high as 5000; Supplemental Dataset 1), the DEG *Sopen07g006810* had moderate levels of expression in medium- and high-acylsugar-producing accessions, and very low levels of expression in low-acylsugar-producing accessions (Figure 2C). Putative orthologs of *Sopen07g006810* and *Sopen08g020180* in *S. lycopersicum (Solyc07g017950* and *Solyc08g066360*, respectively; Supplemental Dataset 3) show 47- and 16-fold higher expression, respectively, in isolated stem trichomes compared to shaved stems (Ning et al., 2015).

Two other genes *(Sopen03g040490* and *Sopen01g042050)* annotated as ‘invertase’ and ‘sugar transporter ERD6-like isoform 5’, respectively, were also upregulated in high-acylsugar-producing accessions (Table 3). ERD6-like transporters are capable of monosaccharide uptake by facilitated diffusion (Yamada et al., 2010). The putative ortholog of *Sopen03g040490* (invertase) in S. *lycopersicum (Solyc03g121680)* shows no expression in isolated trichomes, whereas *Solyc01g098490* (putative ortholog of the monosaccharide transporter *Sopen01g042050;* Supplemental Dataset 3) shows 22-fold higher expression in trichomes compared to non-trichome tissues (Ning et al., 2015).

### Genes putatively encoding oxylipin metabolic proteins and other defense proteins are upregulated in low-acylsugar-producing accessions

Because acylsugars have important roles in plant defense (Hawthorne et al., 1992; Juvik et al., 1994; Liedl et al., 1995; Chortyk et al., 1997; Hare, 2005; Luu et al., 2017), we investigated if defense genes were differentially expressed between low- and high-acylsugar-producing accessions. Nine genes putatively encoding fatty acid desaturase and lipoxygenase enzymes were differentially expressed, and eight of them were upregulated in low-acylsugar-producing accessions (Table 4). Lipoxygenases catalyze the dioxygenation of polyunsaturated fatty acids produced by fatty acid desaturases, and this enzymatic conversion is the first step in the biosynthesis of oxylipins, which contribute to innate immunity (La Camera et al., 2004). Oxylipin response pathways are regulated by TGA transcription factors, and two classes of enzyme (oxo-phytodienoate reductase and glutathione S-transferase) have been reported to take important roles in metabolizing highly reactive oxylipins into less reactive molecules (Mueller et al., 2008). Genes putatively encoding these oxylipin signaling and metabolic proteins were also upregulated in low-acylsugar-producing accessions (Table 4).

Lipoxygenase-produced oxylipins have important roles in plant defense against herbivores (Halitschke and Baldwin, 2003) and microbial pathogens (Prost et al., 2005). Therefore, low-acylsugar-producing accessions may generate oxylipins to compensate for diminished antiherbivory and antimicrobial activities of acylsugars. Consistent with this hypothesis, genes putatively encoding subtilisin-like proteases, which play important role in the recognition of plant pathogens and activation of immune responses (Figueiredo et al., 2014), and additional proteins involved in plant defense response to fungal and other biotic stimulus were also upregulated in low-acylsugar-producing accessions (Table 4). Based on expression profiles, genes involved in fatty acid desaturation, lipoxygenation, oxylipin signaling and metabolism, and antimicrobial defense showed moderate to strong correlation with each other (Supplemental Figure 6), indicating an active network of innate immunity in low-acylsugar-producing accessions.

**Figure 6.**
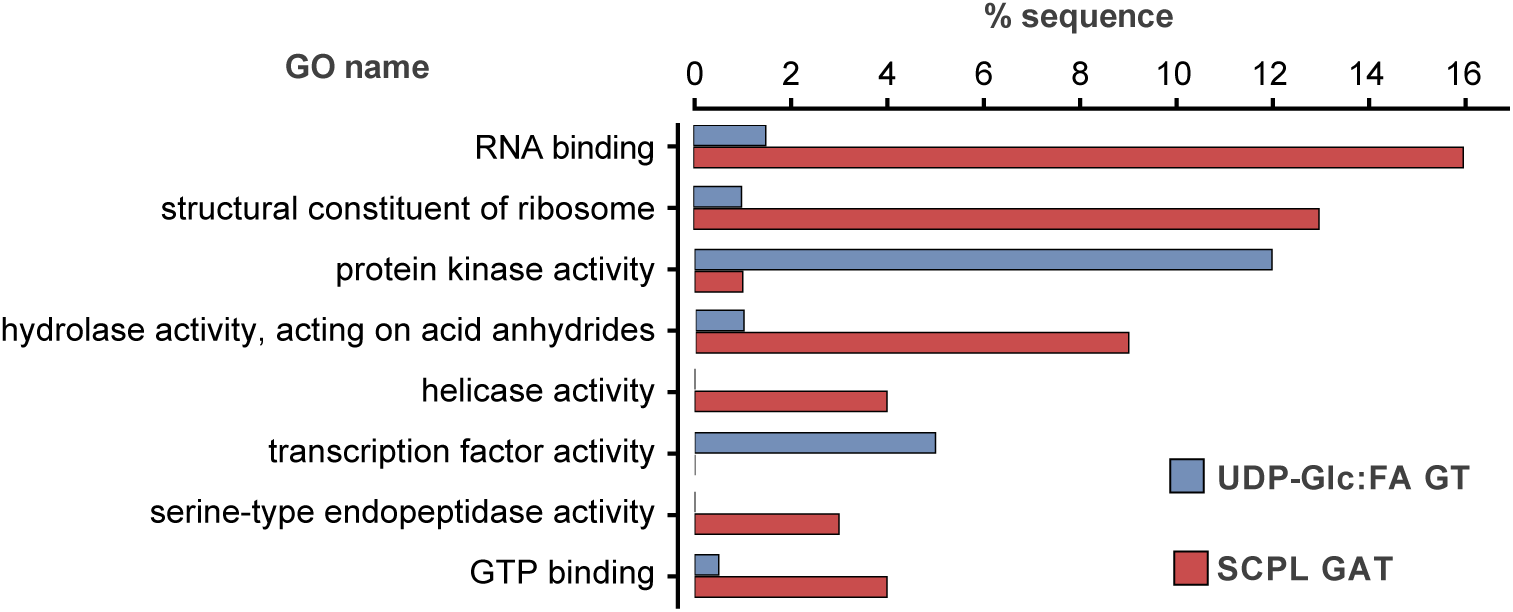
Enrichment analysis of Gene Ontology (GO) terms associated with genes most strongly correlated with two genes encoding known acylglucose biosynthetic enzymes (UDP-Glc:FA GT and SCPL GAT). Enrichment analysis was performed with Blast2GO (Conesa et al., 2005) using Fischer’s exact test (P < 0.05). Top-100 genes associated with *Sopen01g049990* (UDP-Glc:FA GT) in both ‘lightcyan’ and ‘tan’ modules, and top-100 genes associated with *Sopen10g020280* (SCPL GAT) in ‘green’ module were used for the analysis. Categories are listed from top to bottom based on their P values (from P= 3.57E-06 for ‘RNA binding’ to P= 4.39E-02 for ‘GTP binding’). GO terms in ‘molecular function’ subsection only were selected, and redundant categories were combined. A full list of GO terms and associated P values are given in Supplemental Figure 8.

### Amine oxidase-related defense response genes are upregulated in high-acylsugar-producing accessions

We found that one DEG (*Sopen06g005510*) predicted to encode the catalytic domain of glutamine synthetase had 183-fold higher expression in the ‘HIGH’ group than in the ‘LOW’ group (FDR= 2.60E-42). In plants, expression of glutamine synthetase is induced by higher concentrations of ammonia, which encouraged us to search for DEGs involved in ammonia-liberating reactions. Three genes encoding amine oxidase domains *(Sopen07g017370, Sopen11g004040*, and *Sopen11g004050)* and an aldehyde oxidase gene *(Sopen01g035050)*, whose product could potentially oxidize aldehydes produced by the amine oxidases, were also noticeably upregulated in high-acylsugar-producing accessions (Table 4). BLAST results indicate that *Sopen11g004040* and *Sopen11g004050* show similarity with FLOWERING LOCUS D-like proteins, which have a flavin-containing amine oxidase domain and are required for systemic acquired resistance in *Arabidopsis thaliana* (Singh et al., 2013).

H_2_O_2_ generated by amine oxidase- and aldehyde oxidase-catalyzed reactions acts as a signal molecule and plays an important role during pathogen invasion by acting on extensin proteins (Niebel et al., 1993; Jackson et al., 2001; Cona et al., 2006). Two genes predicted to encode extensin-like glycoproteins were upregulated in high-acylsugar-producing accessions (Table 4), and one of them *(Sopen05g003410)* showed strong positive correlation with amine oxidase- and aldehyde oxidase-encoding genes (Supplemental Figure 6). Although amine oxidase-related defense genes were upregulated in high-acylsugar-producing accessions, it is important to note that their expression levels were much lower than some oxylipin-related defense genes in low-acylsugar-producing accessions (Figure 4).

### Identification of co-expressed gene networks associated with acylsugar metabolism and plant defense

Strong correlation among expression profiles of known and putative acylsugar metabolic genes (Figure 3) encouraged us to perform gene co-expression network analysis. Using weighted gene correlation network analysis (WGCNA) (Langfelder and Horvath, 2008), we identified 15 modules (clusters of co-expressed genes). As a representative of gene expression profile in each WGCNA module, a module eigengene was defined as the first principle component in each module (Langfelder and Horvath, 2008). Acylsugar accumulation and ASAT gene expression showed strong positive correlation with module eigengene of the ‘turquoise’ module, and strong negative correlation with module eigengene of the ‘black’ module (Figure 5). Of the 586 upregulated genes in high-acylsugar-producing accessions, 545 (93%) were found in the ‘turquoise’ module, whereas 371 (74%) of the 501 downregulated genes were clustered in the ‘black’ module (Supplemental Dataset 4: Sheet 1). The top-100 genes most strongly correlated with acylsugar accumulation in the ‘turquoise’ module included genes involved in BCAA/BCFA metabolism, SCFA metabolism (putative FAS components and acyl-activating enzymes AAE1), acylsucrose metabolism (except *Sp-ASH1)*, transport process (ABC transporters), and central carbon metabolism (Supplemental Dataset 4: Sheet 3). On the other hand, the ‘black’ module contained genes for putative oxylipin-mediated defense pathway, and two carboxylesterase/acylhydrolase genes *(Sopen02g020660* and *Sopen02g020670)* that exhibited strong negative correlation with acylsugar accumulation (*ρ=* -0.79 and -0.85, respectively; *P* = 4.56E-07 and 3.84E-09, respectively).

We also selected *Sopen01g002520* (lipoxygenase) and *Sopen11g004040* (FLOWERING LOCUS D) as representative of the oxylipin-mediated and the amine oxidase-mediated defense networks, respectively, and identified genes that are strongly correlated with them. Genes putatively encoding fatty acid desaturase, TGA transcription factor, glutathione S-transferase, subtilisin-like protease and other pathogenesis-related proteins were associated with the lipoxygenase gene (Supplemental Dataset 4: Sheet 7). On the other hand, a search for top-100 genes associated with *Sopen11g004040* in the ‘turquoise’ module identified the adjacent FLOWERING LOCUS D gene *Sopen11g004050*, as well as genes predicted to encode copper-containing amine oxidase, aldehyde oxidase, extensin-like protein, leucine rich repeat (LRR) containing serine/threonine protein kinase, tyrosine protein kinase, NB-ARC domain-containing protein, exostosin family glycosyltransferases, F-box proteins, transcription factors (Supplemental Table 4), and more than 50 hypothetical proteins (Supplemental Dataset 4: Sheet 8).

It is interesting to note that a protamine P1-like protein (Sopen09g015380; 154-aa; 24 Lys, 29 Arg) and some hypothetical proteins encoded by the genes strongly correlated with *Sopen11g004040*, had higher content of Lys and/or Arg residues (Supplemental Figure 7). The biological implication of this observation is not clear; however, small, cationic peptides from other plants have antimicrobial properties (Marcos et al., 2008), and co-expression of these genes with *Sopen11g004040* may reflect another branch of the plant defense network.

**Figure 7.**
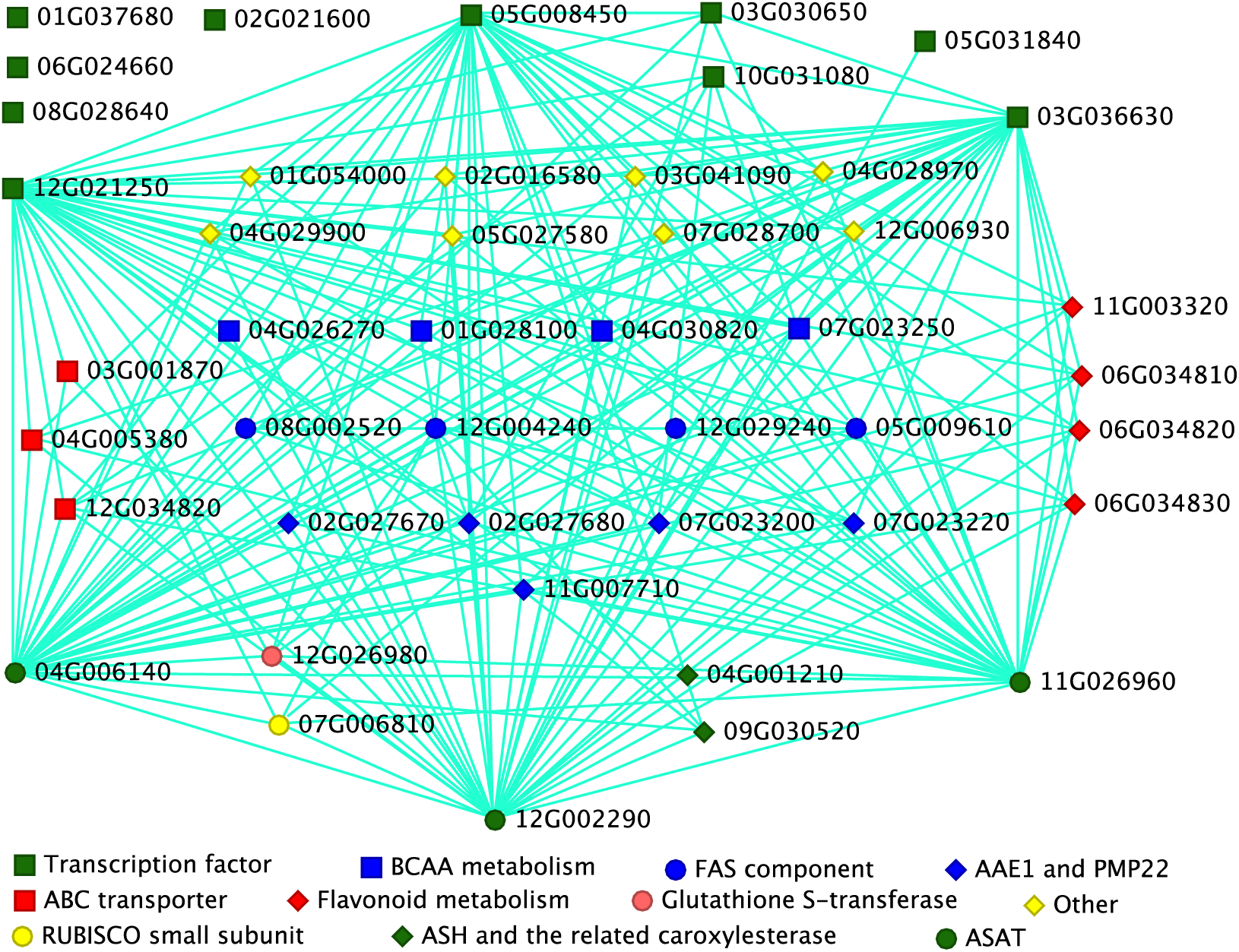
Simplified acylsugar metabolic gene network. Nodes and edges represent genes and intramodular connectivities, respectively. We used WGCNA to identify the top-50 genes most strongly connected to each of the eight transcription factor genes (all green squares except *Sopen03g030650* and *Sopen05g031840*) that responded to imazapyr and were in the ‘turquoise’ module, as well as three acylsucrose biosynthetic genes (three ASATs). These 11 top-50 lists contained only 160 different genes. Using VisANT (Hu et al., 2005), with an edge weight cutoff value of 0.20, 41 genes were identified as most strongly connected. Three genes putatively encoding AP2-family transcription factors *(Sopen05g008450, Sopen12g021250*, and *Sopen03g036630)* showed strongest connectivity with acylsugar (and also flavonoid) metabolic genes. This network also includes two additional transcription factor genes *(Sopen03g030650* and *Sopen05g031840). Sopen03g030650* was downregulated only at 1 mM imazapyr treatment, whereas *Sopen05g031840* was slightly upregulated at both 0.1 and 1 mM imazapyr treatment.

### Protein kinases are correlated with the UDP-Glc:FA GT, whereas factors involved in translation are strongly associated with the SCPL GAT

Because expression profiles of the two phase 2 acylglucose biosynthetic genes were not consistent with acylsugar accumulation (Table 2), we attempted to find and analyze genes that are strongly associated with them. The expression profile of *Sopen01g049990* (UDP-Glc:FA GT) showed strong correlation with module eigengenes in the ‘lightcyan’ and ‘tan’ modules, whereas *Sopen10g020280* (SCPL GAT) was associated with the ‘green’ module (Figure 5). We selected top-100 genes associated with each of *Sopen01g049990* and *Sopen10g020280* in these modules, and enrichment analysis revealed that Gene Ontology (GO) terms such as ‘phosphotransferase activity’ (GO:0016773; *P=* 3.32E-04), ‘protein kinase activity’ (GO:0004672; *P=* 5.82E-04), and ‘transcription factor activity’ (GO:0003735; P= 3.40E-02) were overrepresented in the ‘lightcyan’ and ‘tan’ modules; on the other hand, GO terms such as ‘RNA binding’ (GO:0003723; P= 3.57E-06) and ‘structural constituent of ribosome’ (GO:0003735; P= 1.96E-05) were significantly overrepresented in the ‘green’ module (Figure 6; Supplemental Figure 8). In fact, 11 genes encoding ribosomal proteins, as well as many genes involved in post-transcriptional processing and translation process were in the list of top-100 genes associated with *Sopen10g020280* (Supplemental Table 5). Interestingly, this list also included two genes encoding signal recognition particles, consistent with a previous report that the SCPL GAT is a glycoprotein with a cleavable N-terminal 18-aa signal peptide (Li and Steffens, 2000).

**Table 5.** Response of selected genes to imazapyr treatment.

| Gene ID | Imazapyr 0.1 mM |  | Imazapyr 1 mM |  | Annotation |
| --- | --- | --- | --- | --- | --- |
|  | Log<sub>2</sub>FC | FDR | Log<sub>2</sub>FC | FDR |  |
| Genes related to phase 1 of acylsugar biosynthesis (Table 1 genes) |  |  |  |  |  |
| Sopen11g004560 | 0.40 | 3.35E-01 | -2.22 | 8.69E-12 | Acetolactate synthase |
| Sopen07g027240 | 0.92 | 3.20E-03 | 1.00 | 2.95E-04 | Ketol-acid reductoisomerase |
| Sopen05g032060 | -0.08 | 8.46E-01 | -1.22 | 1.32E-06 | Dihydroxy-acid dehydratase |
| Sopen08g005060 | 2.03 | 1.45E-10 | 1.91 | 1.68E-10 | Isopropylmalate synthase |
| Sopen08g005140 | -3.11 | 6.11E-06 | -5.63 | 7.20E-12 | Isopropylmalate synthase |
| Sopen04g030820 | -2.90 | 5.88E-08 | -3.73 | 1.11E-12 | Branched-chain aminotransferase-2 |
| Sopen04g026270 | -0.70 | 7.42E-02 | 3.17 | 1.18E-21 | Branched-chain keto acid dehydrogenase E1 subunit |
| Sopen01g028100 | -0.72 | 2.61E-02 | 1.91 | 1.35E-12 | Branched-chain keto acid dehydrogenase E2 subunit |
| Sopen07g023250 | -2.65 | 2.50E-05 | -7.04 | 1.92E-19 | 3-Hydroxyisobutyryl-CoA hydrolase |
| Sopen05g023470 | -0.68 | 3.24E-01 | -1.41 | 8.99E-03 | 3-Hdroxyisobutyryl-CoA hydrolase |
| Sopen12g032690 | -1.28 | 4.23E-02 | -4.52 | 9.34E-10 | Mitochondrial acyl-CoA thioesterase |
| Sopen12g004240 | -1.55 | 1.88E-06 | -2.50 | 8.84E-16 | KAS IV/ KAS II-like domain |
| Sopen08g002520 | -2.29 | 9.29E-08 | -4.03 | 7.27E-20 | Beta-ketoacyl-ACP synthase III |
| Sopen05g009610 | -2.81 | 3.03E-05 | -4.89 | 1.08E-11 | Beta-ketoacyl-ACP reductase |
| Sopen12g029240 | -2.25 | 5.40E-05 | -3.91 | 4.51E-12 | Enoyl-ACP reductase domain |
| Sopen02g027670 | -2.29 | 2.18E-04 | -4.73 | 6.94E-13 | Acyl-activating enzyme 1 |
| Sopen02g027680 | -1.83 | 2.71E-04 | -4.95 | 1.78E-19 | Acyl-activating enzyme 1 |
| Sopen07g023200 | -2.28 | 4.84E-05 | -4.86 | 1.15E-14 | Acyl-activating enzyme 1 |
| Sopen07g023220 | -2.64 | 2.13E-06 | -6.08 | 4.25E-20 | Acyl-activating enzyme 1 |
| Genes related to phase 2 of acylsugar biosynthesis (Table 2 genes) |  |  |  |  |  |
| Sopen01g049990 | 1.81 | 2.59E-06 | 1.81 | 4.01E-07 | UDP-glucose:fatty acid glucosyltransferase |
| Sopen10g020280 | 0.83 | 4.02E-03 | 0.06 | 8.40E-01 | Serine carboxypeptidase-like glucose acyltransferase |
| Sopen12g002290 | -3.07 | 7.59E-06 | -6.55 | 1.65E-14 | Acylsugar acyltransferase 1 |
| Sopen04g006140 | -1.62 | 1.23E-02 | -5.99 | 2.64E-15 | Acylsugar acyltransferase 2 |
| Sopen11g026960 | -1.40 | 1.69E-03 | -4.30 | 6.51E-22 | Acylsugar acyltransferase 3 |
| Sopen05g030120 | -1.16 | 1.33E-04 | -1.91 | 4.26E-12 | Acylsugar acylhydrolase 1 |
| Sopen05g030130 | -1.40 | 3.68E-04 | -4.53 | 6.52E-29 | Acylsugar acylhydrolase 2 |
| Sopen09g030520 | -1.05 | 9.93E-04 | -1.81 | 4.01E-10 | Acylsugar acylhydrolase 3 |
| Sopen04g001210 | -2.34 | 5.74E-11 | -6.12 | 8.82E-50 | Carboxylesterase |
| Other genes related to acylsugar metabolism (Table 3 genes) |  |  |  |  |  |
| Sopen04g023150 | -0.013 | 9.64E-01 | -0.24 | 5.11E-01 | ABC transporter F family member |
| Sopen01g047950 | 0.39 | 5.51E-01 | 1.26 | 6.38E-03 | ABC transporter G family member |
| Sopen04g005380 | -1.19 | 3.17E-03 | -2.71 | 1.05E-12 | ABC transporter G family |
| Sopen12g034820 | -2.46 | 1.74E-05 | -5.01 | 4.43E-14 | Pleiotropic drug resistance protein 1-like (ABC-G family) |
| Sopen03g001870 | -1.89 | 3.65E-05 | -2.81 | 1.34E-10 | ABC transporter B family |
| Sopen07g006810 | -3.25 | 1.92E-05 | -5.50 | 4.99E-10 | RUBISCO small subunit |
| Sopen08g020180 | -1.42 | 3.69E-05 | -3.33 | 3.41E-23 | NADP-dependent malic enzyme |
| Sopen03g040490 | -1.32 | 3.35E-04 | -3.67 | 1.96E-23 | Invertase (beta-fructofuranosidase) |
| Sopen01g042050 | -2.17 | 7.50E-03 | -2.43 | 6.64E-04 | Sugar transporter ERD6-like |
| Sopen05g008450 | -1.58 | 8.73E-03 | -4.58 | 2.43E-12 | AP2 domain |
| Sopen12g021250 | -2.87 | 1.93E-04 | -1.56 | 1.74E-02 | AP2 domain |
| Sopen03g036630 | -1.89 | 5.15E-04 | -2.95 | 1.71E-08 | AP2 domain |
| Sopen10g031080 | -1.19 | 3.82E-02 | -2.31 | 8.15E-06 | Homeobox associated leucine zipper |
| Sopen06g024660 | -1.22 | 6.20E-03 | -2.32 | 1.46E-08 | Myb-like DNA-binding domain |
| Sopen08g028640 | -1.59 | 1.02E-02 | -2.31 | 5.08E-05 | AP2 domain |
| Sopen02g021600 | -1.24 | 4.25E-03 | -1.80 | 4.45E-06 | Myb-like DNA-binding domain |
| Sopen01g037680 | -1.46 | 3.33E-02 | -3.52 | 1.82E-07 | TCP family transcription factor |
| Sopen03g037210 | -1.88 | 8.94E-04 | -2.49 | 1.33E-06 | Helix-loop-helix DNA-binding domain |

- 1 Log<sub>2</sub>FC and FDR indicate log<sub>2</sub> (fold-change) and false discovery rate (*P* values adjusted for - 2 multiple-testing), respectively. Positive and negative log<sub>2</sub>FC values indicate higher and lower - 3 expression levels, respectively in response to the inhibitor treatment compared to control - 4 solution.

**Figure 8.**
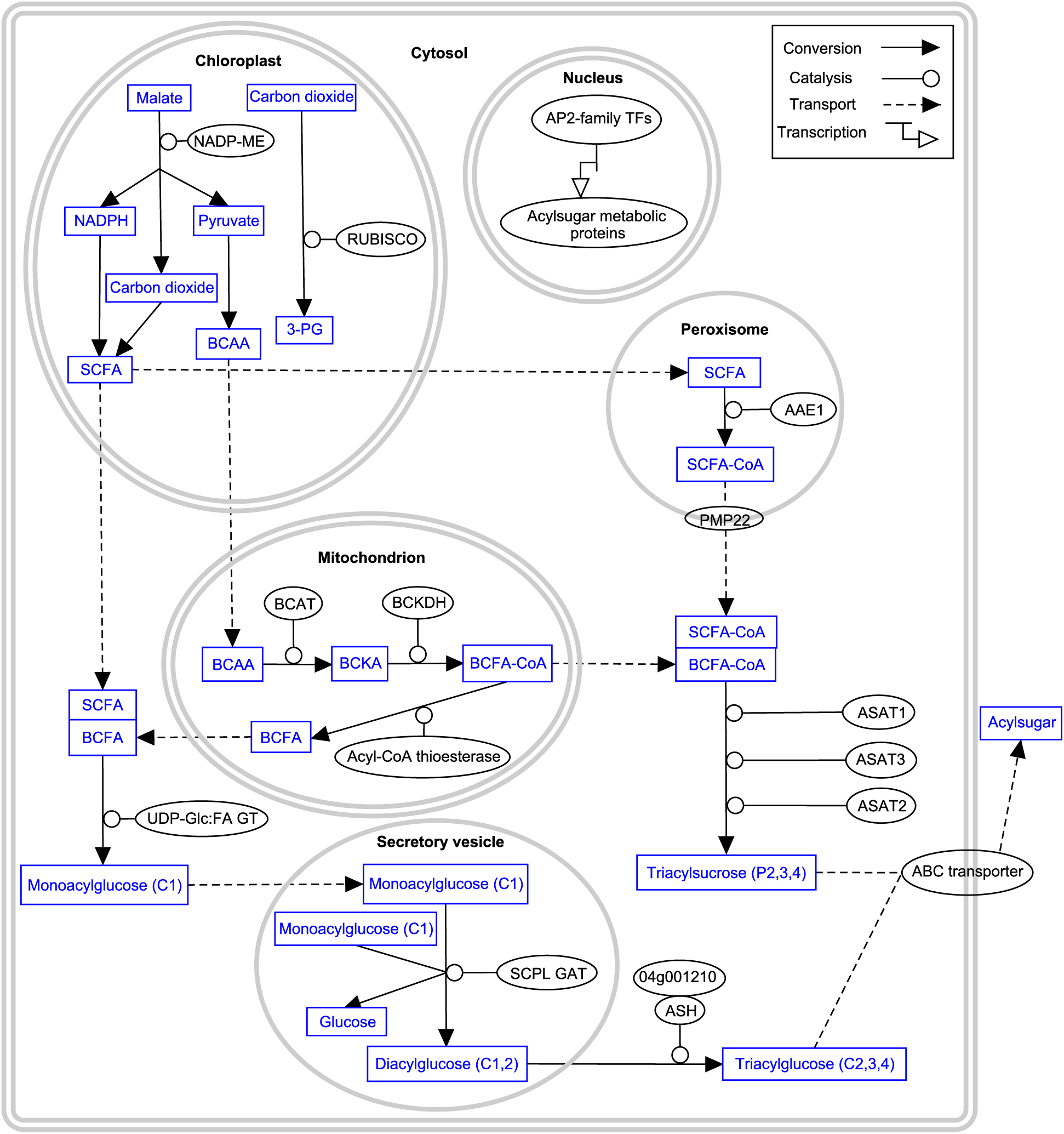
Proposed model of acylsugar metabolism in *S. pennellii.* Role of different subcellular compartments are shown. Metabolites are highlighted in blue within boxes. Enzymes, transporters and transcription factors (TFs) with known and putative functions in acylsugar metabolism are highlighted in black within ovals. Single arrow does not necessarily represent a single enzymatic step. This figure was prepared using WikiPathways (Pico et al., 2008). Abbreviations: NADP:ME= NADP-malic enzyme; 3-PG= 3-phosphoglycerate; BCKA= branched-chain keto acid.

Because *Sopen10g020280* (SCPL GAT) does not show significant differential expression between low- and high-acylsugar-producing accessions, we considered the possibility that genes associated with it in the ‘green’ module were simply housekeeping genes whose expression did not vary, either. However, 9 of our 10 selected housekeeping genes, as well as many genes putatively encoding signal recognition particles and ribosomal proteins, showed little or no correlation with *Sopen10g020280* and were placed in other modules (Supplemental Table 6). These results suggest that some genes in the ‘green’ module, whose expression profiles are strongly correlated with that of *Sopen10g020280*, may have a role in regulating the SCPL GAT activity at the level of translation.

### Inhibition of BCAA biosynthesis changes transcript levels of many acylsugar candidate DEGs, but not defense DEGs

Acetolactate synthase (ALS) catalyzes the first common step in the biosynthesis of BCAAs (Supplemental Figure 4A), and inhibition of this enzyme significantly lowers acylsugar production in *S. pennellii* (Walters and Steffens, 1990). To determine whether inhibition of this enzyme changed expression of ALS and other candidate genes involved in acylsugar metabolism, we treated leaves of the high-acylsugar-producing accession LA0716 with the ALS inhibitor imazapyr at 0.1 mM and 1 mM concentrations. Many of the DEGs involved in BCAA/BCFA metabolism showed differential expression in response to imazapyr treatment (Table 5). Interestingly, at 1 mM imazapyr, expression of *Sopen04g026270* and *Sopen01g028100* (branched-chain keto acid dehydrogenase complex E1 and E2 subunits, respectively) was significantly increased, possibly because the amount of 2-ketobutyrate, a substrate of ALS that also can be used directly by the dehydrogenase complex to make propionyl-CoA, builds up (Walters and Steffens, 1990).

Surprisingly, DEGs putatively involved in SCFA metabolism (FAS components and AAE1 members), showed significant downregulation at both 0.1 mM and 1 mM imazypyr in a concentration-dependent manner. It is interesting to note that AAE1 proteins have a peroxisomal location in *Arabidopsis thaliana* (Reumann et al., 2009); one gene *(Sopen11g007710)* predicted to encode Mpv17/PMP22 family peroxisomal membrane protein had 7.7-fold (FDR= 1.17E-08) and 165-fold (FDR= 3.62E-24) lower expression at 0.1 mM and 1 mM imazapyr treatment, respectively (Supplemental Dataset 5). *Sopen11g007710* was upregulated 29-fold in the ‘HIGH’ group (FDR= 2.80E-11; Supplemental Dataset 1), and its putative ortholog in S. *lycopersicum (Solyc11g012990;* Supplemental Dataset 3) has 1420-fold higher expression in isolated trichomes than shaved stems, consistent with trichome-enriched expression of AAE1 members (Ning et al., 2015).

*Sopen01g049990* (UDP-Glc:FA GT; phase 2 of acylglucose biosynthesis) showed 3.5-fold higher expression at both concentrations of imazapyr (FDR= 2.59E-06 and 4.01E-07, respectively), supporting our previous finding that a negative correlation may exist between acylsugar level and expression level of this gene. *Sopen10g020280* (SCPL GAT) did not show any significant change in expression level at either concentration (log_2_FC > 1, FDR < 0.05). On the other hand, DEGs encoding ASATs (phase 2 of acylsucrose biosynthesis) showed significant decrease in gene expression level in a concentration-dependent manner (by as much as 94-fold, FDR= 1.65E-14), as did ASHs and the related carboxylesterase *Sopen04g001210* (by as much as 70-fold, FDR= 8.82E-50) (Table 5).

Three DEGs putatively encoding ABC transporters were significantly downregulated at both concentrations of imazapyr (by as much as 32-fold, FDR= 4.43E-14). Imazapyr treatment in *Arabidopsis thaliana* resulted in significant induction of nine genes (out of the ten genes reported) encoding ABC transporters, indicating their role in detoxification process (Manabe et al., 2007). Similarly, 15 putative ABC transporter genes in *S. pennellii* were upregulated in response to imazapyr either at higher concentration or at both concentrations (Supplemental Table 7). However, three putative ABC transporter DEGs were repressed by imazapyr treatment, consistent with their inclusion in the WGCNA ‘turquoise’ module with other acylsugar metabolic genes.

Interestingly, DEGs involved in central carbon metabolism were also significantly downregulated in response to imazapyr (by as much as 45-fold, FDR= 4.99E-10 for RUBISCO small subunit; Table 5). While investigating the role of central carbon metabolism in specialized metabolism of *S. lycopersicum* and *S. habrochaites* glandular trichomes, Balcke et al. (2017) reported importance of glutathione metabolism to cope with oxidative stress associated with high productivity of specialized metabolites in trichomes (Balcke et al., 2017). One glutathione S-transferase gene (*Sopen12g026980*) was upregulated in high-acylsugar-producing accessions (Figure 4) and was placed in the ‘turquoise’ module due to strong positive correlation between expression profile of this gene and acylsugar accumulation (ρ= 0.82; P= 5.93E-08). *Sopen12g026980* showed 2- and 18-fold (FDR= 2.36E-02 and 8.94E-18, respectively) downregulation at 0.1 mM and 1 mM imazapyr, respectively (Supplemental Dataset 5). According to data of Ning et al. (2015), the putative ortholog of this gene in S. *lycopersicum (Solyc05g006730)* shows 287-fold higher expression in isolated stem trichomes compared to underlying tissues.

The 14 genes strongly associated with the FLOWERING LOCUS D gene *Sopen11g004040* (Supplemental Table 4) did not show any significant differential expression (log_2_FC > 1, FDR < 0.05) in response to imazapyr treatment at either concentration, except two genes which showed slight changes in gene expression only at one concentration (Supplemental Table 8). On the other hand, DEGs with known and putative functions in acylsugar metabolism responded considerably to ALS inhibition. These results suggested that additional genes involved in acylsugar metabolism may also be represented in the list of 171 DEGs (out of 1087 common DEGs between low- and high-acylsugar-producing accessions) that responded to imazapyr at both concentrations (Supplemental Dataset 5). In this list, 13 genes putatively encoding transcription factors were present; of these, nine had higher expression levels in high-acylsugar-producing accessions (Table 3), and eight of these were found in the WGCNA ‘turquoise’ module.

### Three AP2-family transcription factor genes form a strong co-expression network with many acylsugar and flavonoid metabolic genes

We selected the top-50 most strongly connected genes (based on intramodular connectivity as determined by WGCNA) for each of the eight above-mentioned transcription factor genes, and merged them to identify the most strongly interconnected gene network. Three genes *(Sopen12g021250, Sopen03g036630*, and *Sopen05g008450*) putatively encoding AP2-family transcription factors and one gene *(Sopen10g031080)* encoding a homeobox-associated leucine-zipper protein revealed strongest intramodular connectivity with acylsugar metabolic genes (Figure 7). Astonishingly, 48 genes were common among the top-50 most strongly connected genes for each of the three AP2-family members (Supplemental Figure 9).

This network included genes involved in BCFA synthesis, SCFA synthesis (FAS components) and their activation (AAEs and the related peroxisomal membrane protein Mpv17/PMP22), acylsucrose metabolism (three ASATs, ASH3 and the related carboxylesterase *Sopen04g001210)*, as well as genes putatively encoding three ABC transporters, RUBISCO small subunit, glutathione S-transferase, and other proteins. According to data of Ning et al. (2015), putative orthologs of the three AP2-family transcription factor genes in S. *lycopersicum (Solyc12g042210, Solyc03g117720*, and *Solyc05g013540*, respectively; Supplemental Dataset 3) show more than 205-, 24-, and 252-fold higher expression in isolated trichomes compared to shaved stems, whereas *Solyc10g080540* (putative ortholog of *Sopen10g031080;* leucine-zipper protein) does not show any significant difference in expression between isolated trichomes and shaved stems (1.2-fold, FDR= 1).

However, not all genes in this network had an obvious connection to acylsugar metabolism. One gene on chromosome 11 (*Sopen11g003320;* UDP-glucose:catechin glucosyltransferase (Noguchi et al., 2008)) and three sequential genes on chromosome 6 *(Sopen06g034810, Sopen06g034820*, and *Sopen06g034830;* myricetin methyltransferase (Schmidt et al., 2012; Kim et al., 2014)) showed strong intramodular connectivity with AP2-family transcription factors and other acylsugar metabolic genes (Figure 7). These flavonoid metabolic genes were also strongly downregulated in response to imazapyr treatment (Supplemental Figure 10), and their putative orthologs in *S. lycopersicum* are preferentially expressed in trichomes (Ning et al., 2015). In addition to abundant acylsugar compounds, presence of flavonoid compounds, such as methylated myricetin, has been reported in S. *pennellii* (LA0716) trichomes (McDowell et al., 2011). These results suggest that a subset of AP2-family transcription factors is strongly connected with metabolism of both acylsugar and flavonoid compounds.

## DISCUSSION

Acylsugars are powerful natural pesticides (Puterka et al., 2003), and increasing acylsugar-mediated insect pest resistance has been a target of various breeding programs in solanaceous crops (Bonierbale et al., 1994; Lawson et al., 1997; Alba et al., 2009; Leckie et al., 2012). The success of these breeding programs and bioengineering of acylsugar production will largely depend on unraveling the complex network of structural and regulatory genes. Here, we exploited natural variation in acylsugar production among *S. pennellii* accessions through a combination of comparative transcriptomics study and gene co-expression network analysis to identify a number of genes with known and putative functions in acylsugar metabolism. Our initial hypothesis for this work was that expression levels of known acylsugar biosynthetic genes would be consistent with acylsugar levels in different accessions of *S. pennellii*, and that additional, unknown genes involved in acylsugar metabolism would be coordinately expressed with them. Our approach was validated by the identification of several DEGs with known functions in acylsugar biosynthesis, such as BCAA/BCFA metabolic genes and acylsucrose biosynthetic genes (ASATs). Our work also revealed differential expression of two defense networks among *S. pennellii* accessions.

### Defense pathway genes

Higher expression levels of genes putatively encoding oxylipin metabolic proteins, subtilisin-like proteases, and other defense proteins in low-acylsugar-producing accessions (Table 4) may indicate a compensatory mechanism for diminished defense activities of acylsugars in these accessions. Alternatively, expression of these defense genes may be suppressed in high-acylsugar-producing accessions due to the protection provided by acylsugars. Our data do not allow us to distinguish between these two possibilities, but it is important to note that all plants used in this study were grown in the same growth chamber, and no insects or symptoms of disease were apparent on any plant.

A different class of defense genes, including those putatively encoding amine oxidases, aldehyde oxidase, and extensin-like proteins, was upregulated in high-acylsugar-producing accessions. In general, these amine oxidase-related DEGs had much lower expression levels than some oxylipin-related defense DEGs in low-acylsugar-producing accessions (Figure 4). These different levels of expression may indicate that defense DEGs in high-acylsugar-producing accessions are involved in baseline innate immunity, whereas acylsugars provide the major defensive barrier. During pathogen or herbivore attack, these amine oxidases may be upregulated to produce H_2_O_2,_ which can activate extensin proteins (Cona et al., 2006). Increased levels of extensin transcripts upon nematode infection has been reported in tobacco (Niebel et al., 1993), and deposition of extensin mediated by H_2_O_2_-induced peroxidase during callus culture elicitation has been reported in *Vitis vinifera* (Jackson et al., 2001).

Most defense DEGs (both oxylipin-related and amine oxidase-related) had similar expression levels in the ‘MEDIUM’ accession LA1302 and in low-acylsugar-producing accessions (Figure 4). Multi-dimensional scaling plots also showed that the overall gene expression profile of LA1302 is similar to that of low-acylsugar-producing accessions, but very different from that of high-acylsugar-producing accessions (Supplemental Figure 2A and 2B). These results are consistent with the previous prediction that LA1302 is more closely related to low-acylsugar-producing accessions than high-acylsugar-producing accessions (Shapiro et al., 1994). Despite the similarity of defense DEGs expression and the overall transcriptome profile between LA1302 and low-acylsugar-producing accessions, many DEGs with known and putative functions in acylsugar metabolism showed intermediate expression levels in LA1302 (Figure 2). Together, these results suggest that there is no strict linear relationship between acylsugar accumulation and expression of defense pathways. Further studies are required to fully understand this interaction.

### Candidate genes involved in acylsugar biosynthesis and export

Strong correlations among expression profiles of selected acylsugar metabolic genes (Figure 3) encouraged us to perform gene co-expression network analysis, which identified candidate genes involved in acylsugar metabolism. Roles of candidate genes were further supported by their response to imazapyr treatment, which caused significant downregulation of most acylsugar metabolic genes (Table 5), but not defense genes (Supplemental Table 8). Additionally, data of Ning et al. (2015) indicate that putative *S. lycopersicum* orthologs of most of our candidate genes are preferentially expressed in trichomes. Roles of some candidate genes in acylsugar metabolism are discussed below.

Acylsugar phase 1: Sequence similarity between Sopen12g004240’s KAS domain and *Cuphea* KAS IV/KAS II-like enzymes, which synthesize medium-chain (C8-C12) SCFAs, suggests a role of Sopen12g004240 (in collaboration with Sopen08g002520; KAS III) in acylsugar SCFA synthesis. This is consistent with the previous prediction that KAS enzyme(s) other than KAS I might be involved in SCFA production in S. *pennellii* acylsugars (Slocombe et al., 2008). Furthermore, *Sopen12g004240* is located near a previously identified SCFA quantitative trait locus on the upper arm of chromosome 12 (Blauth et al., 1999).

We identified putative medium-chain specific AAE1 members (Shockey et al., 2003) that may provide SCFA-CoA molecules for ASATs in acylsucrose biosynthesis, based on not only their strong positive correlation with both FAS components and ASATs (Figure 3), but also trichome-enriched expression patterns of their putative orthologs in S. *lycopersicum* (Ning et al., 2015). Duplicated genes often encode proteins with similar but distinct substrate specificity, and DEGs putatively encoding AAE1 members are located on chromosome 2 and chromosome 7 as members of similar gene clusters in both S. *pennellii* (Bolger et al., 2014a) and S. *lycopersicum* (The Tomato Genome Consortium, 2012). It will be interesting to see if different AAE1 members can activate SCFAs with different chain lengths.

Putative peroxisomal locations for the four AAE1 members (Reumann et al., 2009) are consistent with their strong connection with *Sopen11g007710* (Figure 7), which is predicted to encode an Mpv17/PMP22 family peroxisomal membrane protein. PMP22 is presumably involved in controlling permeability of peroxisomal membrane (Brosius et al., 2002), and *Sopen11g007710* showed strong response to imazapyr treatment, suggesting an important role of peroxisome in acylsugar metabolism (Figure 8).

Acylsugar phase 2: Because no tri- or tetra-acylglucose products are formed by the SCPL GAT (Li et al., 1999; Li and Steffens, 2000), synthesis of the final product 2,3,4-tri-*O*-acylglucose from 1,2-di-*O*-acylglucose requires additional, unidentified enzyme(s) (Figure 1A). If these additional enzymatic activities are not regulated at the transcriptional level, then comparative transcriptomics would not be able to identify them. However, transcriptional regulation may be involved in specific segments of this pathway, and the role of few candidate genes can be discussed here.

Although ASHs remove acyl chains from the sucrose pyranose ring in vitro, several possible in vivo functions have been proposed for ASH activity, including roles in acylsugar remodeling, replacing acyl chains, or catalyzing the reverse reaction (adding acyl chains to acylsugar molecules rather than removing them) (Schilmiller et al., 2016). Genes encoding ASH3 and the related carboxylesterase Sopen04g001210 showed strong associations with other acylsugar metabolic genes (Figure 7) and also responded to imazapyr in a similar way (Table 5). It is possible that these enzymes catalyze in vivo acylation of 1,2-di-*O*-acylglucose at C3 and C4 positions and/or removal or remodeling of the C1 acyl group to finally synthesize 2,3,4-tri-*O*-acylglucose. *Sopen04g001210* is of special interest because it has very high expression levels in our leaf samples (Figure 2B), and its putative ortholog in *S. lycopersicum* shows trichome-enriched expression (Ning et al., 2015; Schilmiller et al., 2016). The enzyme encoded by *Sopen04g001210* did not show any detectable activity against tomato acylsucrose fractions (Schilmiller et al., 2016), but it may have an acylglucose-specific function.

Acylsugar export: It is not known how acylsugars are secreted out of trichome cells, or whether acylglucose and acylsucrose molecules are exported by the same mechanism. The SCPL GAT, which catalyzes the second step in phase 2 of acylglucose biosynthesis, has a cleavable N-terminal 18-aa signal peptide, and acylsugars (or at least acylglucose molecules) have been suggested to be secreted via a secretory vesicle pathway (Li et al., 1999; Li and Steffens, 2000). However, to our knowledge, there is no direct evidence to support this hypothesis, and it remains unknown if the final 2,3,4-tri-*O*-acylglucose molecules are synthesized in secretory vesicles or in the cytosol. Here, we identified three putative ABC transporters as candidates for acylsugar (or at least acylsucrose) transport based on their strong intramodular connectivity with ASAT and acylsugar phase 1 genes (Figure 7).

### Role of central carbon metabolism

NADP-malic enzymes catalyze the oxidative decarboxylation of malate to generate pyruvate (required for BCAA biosynthesis), along with CO_2_ and NADPH, both of which can be used for SCFA biosynthesis or carbon fixation. Expression profiles of two genes putatively encoding RUBISCO small subunit and NADP-malic enzyme showed strong positive correlations with acylsugar accumulation (Supplemental Dataset 4). Putative orthologs of these genes in *S. lycopersicum* show trichome-enriched expression (Ning et al., 2015), consistent with the suggestion that these enzymes provide some carbon in S. *pennellii* trichomes to support acylsugar biosynthesis. Strong positive correlations between acylsugar accumulation and transcript levels of putative invertase (whose putative ortholog is not expressed in isolated *S. lycopersicum* trichomes; Ning et al., 2015) and a monosaccharide transporter is consistent with the proposal that *S*. *pennellii* trichomes import at least some photosynthate from underlying leaf tissues to support acylsugar biosynthesis (Slocombe et al., 2008). Recently, the role of central carbon metabolism in supporting specialized metabolism in glandular trichomes of *S. lycopersicum* and *S. habrochaites* was investigated, and it was concluded that glandular trichomes import most of their fixed carbon from underlying leaf tissues, despite being able to perform some photosynthetic activities (Balcke et al., 2017). A role of glutathione metabolism in alleviating oxidative stress was also reported. Our results (Table 3; Figure 7 for glutathione S-transferase) are consistent with these findings.

### Regulation of acylsugar production

Both comparative transcriptomics and imazapyr treatment studies revealed that transcript levels for the two phase 2 acylglucose biosynthetic enzymes (UDP-Glc:FA GT and SCPL GAT) were not positively correlated with acylsugar levels, and any transcriptional regulation of the UDP-Glc:FA GT is negatively correlated with acylsugar production (Table 2; Table 5). Many biochemical pathways are regulated, at least in part, by a feedback inhibitory mechanism, and end-product inhibition often acts on an early rate-limiting step at the enzymatic level. However, it is interesting to note that a negative correlation exists between acylsugar amount and the UDP-Glc:FA GT transcript level. Overrepresentation of Gene Ontology terms ‘protein kinase activity’ and ‘transcription factor activity’ among genes strongly correlated with the UDP-Glc:FA GT (Figure 6) suggests that the UDP-Glc:FA GT activity is controlled by both post-translational and transcriptional regulation. Association of expression patterns of the SCPL GAT with ribosomal proteins and other components of the translational machinery (Supplemental Figure 8) suggests that its activity may be regulated at the translational level.

On the other hand, expression levels of most genes involved in acyl chain synthesis (phase 1) and phase 2 acylsucrose metabolism were positively correlated with acylsugar levels (Figure 2; Supplemental Dataset 4). WGCNA revealed that three genes putatively encoding AP2-family transcription factors form a strong gene co-expression network with many acylsugar and flavonoid metabolic genes. Downregulation of these transcription factor genes in response to imazapyr treatment parallels significant downregulation of most acylsugar and flavonoid metabolic genes (Table 5; Supplemental Dataset 5). Additionally, putative orthologs of these transcription factor genes in *S. lycopersicum* exhibit trichome-enriched expression, similar to many genes involved in acylsugar and flavonoid metabolism (Ning et al., 2015). Together, these results imply important roles for these transcription factors in specialized metabolic pathways of *S. pennellii* trichomes.

Our results suggest that a majority of the acylsugar metabolic pathway, from acyl chain synthesis to acylsugar secretion, represents a remarkable co-expressed genetic network, which may be under a common regulatory mechanism. However, the roles we have proposed for the candidate genes remain to be confirmed by genetic or biochemical studies. Confirmation of common regulatory factors, such as the AP2-family transcription factors, will be useful in genetic manipulation of acylsugar production.

## METHODS

### Plant materials

Three biological replicates were used for each accession in the ‘LOW’ vs. ‘HIGH’ comparison, and for the ‘MEDIUM’ accession LA1302. Four biological replicates were used for each accession in the LA1920 vs. LA0716 comparison. Seeds of all *S. pennellii* accessions were obtained from the C. M. Rick Tomato Genetics Resource Center (University of California, Davis). Seeds were treated with a solution of 20% commercial bleach (1.2% sodium hypochlorite) for 20 minutes, and then germinated on moistened filter paper. Seedlings were transferred to soil at the cotyledon stage, and plants were grown in a growth chamber (24°C day/ 20°C night temperature, 150 mMol m^−2^ s^−1^ photosynthetically active radiation, 16 h photoperiod, and 75% humidity).

### RNA sequencing (RNA-Seq) library preparation and sequencing

Before RNA extraction, secreted acylsugars were removed from the leaf surface of 10-week-old plants by dipping them in ethanol for 2-3 seconds. Leaves were then immediately frozen in liquid nitrogen and stored at -80°C until further use. Total RNA was extracted from leaves using the RNAqueous Total RNA Isolation kit (ThermoFischer). Genomic DNA was removed using the TURBO DNA-free kit (ThermoFischer). Quality of each RNA sample was analyzed with the Agilent 2200 TapeStation software A01.04 (Agilent Technologies). RNA-Seq libraries were prepared from polyA^+^-selected RNA samples using the Illumina TruSeq RNA Library Preparation kit v2, and then sequenced on the Illumina HiSeq 2500 v4 125x125-bp paired-end sequencing platform (High Output Mode) following manufacturer’s specifications. Sequencing was performed at the Texas A&M Genomics and Bioinformatics service center. Sequence cluster identification, quality prefiltering, base calling and uncertainty assessment were done in real time using Illumina’s HCS v2.2.68 and RTA v1.18.66.3 software with default parameter setting. Sequencer.bcl basecall files were demultiplexed and formatted into .fastq files using bcl2fastq v2.17.1.14 script conFigureBclToFastq.pl. Sequencing reads were submitted to the National Center for Biotechnology Information (NCBI) Sequence Read Archive under accession number SRP136022.

### Quality control and mapping of sequencing reads

Phred quality score distributions of sequencing reads were analyzed with FastQC v0.11.4 (Andrews, 2010). All RNA-Seq libraries had average Phred quality scores of more than 35, indicating greater than 99.97% base call accuracy. To remove all the low-quality, ambiguous ‘N’ nucleotides (Phred quality score= 2, ASCII code= #), the following settings were applied for quality control of RNA-Seq reads using Trimmomatic v0.35 (Bolger et al., 2014b)-ILLUMINACLIP:TruSeq3-PE-2.fa:2:30:10, SLIDINGWINDOW= 1:14, MINLEN= 80. Using these settings, more than 98% of the sequencing reads could be retained, and it ensured that none of the nucleotides in the reads were ambiguous.

Processed reads were mapped to the *S. pennellii* genome v2.0 (LA0716) (Bolger et al., 2014a) using TopHat2 v2.1.0 (Kim et al., 2013). The following parameters were used for the mapping process: -mate-inner-dist= 0, -mate-std-dev= 50, -read-realign-edit-dist= 0, -read-edit-dist= 4, -library-type= fr-unstranded, -read-mismatches= 4, -bowtie-n, -min-anchor-len= 8, - splice-mismatches= 0, -min-intron-length= 50, -max-intron-length= 50000, -max-insertion-length= 3, -max-deletion-length= 3, -max-multihits= 20, -min-segment-intron= 50, -max-segment-intron= 50000, -segment-mismatches= 2, -segment-length= 25. A summary of TopHat2 alignment results are given in Supplemental Table 2. Reads mapped to selected loci of interest were visualized with IGV (Thorvaldsdottir et al., 2013).

### Differential gene expression analyses

Uniquely-mapped reads from TopHat2 were counted for all *S*. *pennellii* annotated genes using HTSeq-Count (Anders et al., 2015). The following parameters were used for the counting process: -f bam, -r name, -s no, -m union, -a 20. These count files were used for identification of differentially expressed genes using edgeR (Robinson et al., 2010). Genes with very low expression levels were filtered out before differential testing. For comparison between the ‘LOW’ and ‘HIGH’ groups, genes with more than 1 count per million (CPM) in at least six samples were retained for analysis after read counts were adjusted for transcript lengths. Library sizes were recalculated after the filtering process, and all the samples had post-filter library sizes of more than 99.70%. Tagwise dispersions were calculated (prior.df= 3), and genes were identified as differentially expressed when *P* values, corrected for multiple testing using Benjamini-Hochberg multiple testing correction (Benjamini and Hochberg, 1995), were less than 0.05 (false discovery rate < 0.05), and had at least 2-fold expression differences. For comparison between LA1920 and LA0716, we used the same pipeline, except genes with more than 1 CPM in at least two samples were retained for analysis, and tagwise dispersions were calculated with prior.df= 9. In this analysis, all the samples showed more than 99.85% post-filter library sizes. Lists of differentially expressed genes as determined by edgeR (Robinson et al., 2010) are given in Supplemental Dataset 1.

### Functional annotation of differentially expressed genes

Transcript sequences of differentially expressed genes were extracted from the annotated gene models, and Blast2GO (Conesa et al., 2005) was used for the functional annotation of these transcript sequences. Sequences were searched (BLASTx; word size=3, HSP length cutoff= 33) against the NCBI nonredundant database (subset Viridiplantae, taxa: 33090) with e-value ≤ 1.0E-10, and top-20 BLAST hits were reported. BLAST hits for each sequence were then mapped with the Gene Ontology (GO) terms. Next, annotation was performed with the default parameter settings except the e-value threshold (e-value ≤ 1.0E-10). This process also generated the Enzyme Code (EC) numbers from the Kyoto Encyclopedia of Genes and Genomes (KEGG) pathway. After executing the whole functional annotation process (BLASTx, mapping, and annotation), transcript sequences were exported with sequence description and GO terms. Descriptions of transcript sequences were generated based on similarity levels (e-value and percentage of identity) to the subject genes, as determined by Blast2GO (Conesa et al., 2005).

### Identification of putative orthologs and their trichome-enriched expression

We used reciprocal BLAST to identify putative orthologs of *S. pennellii* genes in *S. lycopersicum.* An all-versus-all BLAST was performed between annotated proteins of *S. pennellii* v2.0 (Bolger et al., 2014a) and *S. lycopersicum* ITAG2.3 (The Tomoto Genome Consortium, 2012; Fernandez-Pozo et al., 2015) with the following parameters-minimum percentage identity=70, minimum percentage query coverage= 50. Pairs with lowest e-value for each BLASTp comparison were selected as putative orthologs. Gene expression profiles of *S. pennellii* putative orthologs in *S. lycopersicum* were obtained from Ning et al. (2015).

### Gene co-expression network analysis

Simple correlation analysis among expression profiles (FPKM values in our 29 samples) of selected genes was performed using the ‘cor’ function in R programming language (R Core Team, 2014). Gene co-expression network analyses and identification of modules (clusters of strongly correlated genes) were performed using the R package WGCNA (Langfelder and Horvath, 2008). Normalized gene expression data (FPKM values) were used as the input, and the 19,378 genes which passed our filtering criteria for minimum expression levels during ‘LOW’ vs. ‘HIGH’ differential testing (*Sopen12g004230* and *Sopen12g004240* were merged) were selected for WGCNA analysis. A matrix of pairwise Spearman’s rank correlation coefficients (SRCCs) between these 19,378 genes was created and then transformed into an adjacency (connection strength) matrix using ‘signed’ network type and a soft threshold (β) value of 20. The value of β was determined based on the scale-free topology fit index (R^2^ > 0.85) and low mean connectivity. The weighted adjacency matrix was then converted to a topological overlap matrix (TOM), and genes were hierarchically clustered based on dissimilarity (dissTOM= 1 - TOM) of the topological overlap. Co-expression modules were defined as the branches of the clustering tree (dendrogram) using the following parameters in ‘cutreeDynamic’ function: distM= disTOM, deepSplit=2, minClusterSize= 200. A module eigengene (ME; representative of gene expression profile in each module) was defined as the first principle component in each module. Modules with very similar expression profiles as determined by their module eigengenes were merged (MEDissThres= 0.2), and a final set of 15 modules were obtained. For each gene, a ‘module membership’ was calculated as the correlation (SRCC) between expression profile of that gene and module eigengene. Associations between external traits and modules were measured as determined by SRCC between trait data and module eigengenes; ‘gene significance’ values for each gene in a module of interest were calculated as the correlation (SRCC) between expression profile of a gene and external trait data. The top-50 intramodular connections for selected genes were visualized by VisANT (Hu et al., 2005).

### Inhibitor studies

Compound leaves of *S. pennellii* LA0716 bearing five leaflets were used for the inhibitor study. Imazapyr (Sigma-Aldrich, USA) was dissolved in distilled water, and administered as 0.1 mM and 1 mM solutions for 36 hours. Cuttings were transferred to small trays containing imazapyr or control solutions, and placed under growth chamber conditions. Total RNA was extracted as described before, and RNA-Seq libraries were prepared using Illumina TruSeq Stranded RNA LT kit. Libraries were sequenced on the Illumina HiSeq 4000 150x150 bp paired-end sequencing platform at the Texas A&M Genomics and Bioinformatics service center, College Station. Raw reads were processed with Trimmomatic (Bolger et al., 2014b), and read-mapping to S. *pennellii* genome was performed with Tophat2 (Kim et al., 2013) using the same parameters as described before, except with the following change: -library-type= fr-firststrand. Uniquely-mapped reads from TopHat2 were counted with HTSeq-Count (Anders et al., 2015) using the following options: -f bam, -r name, -s reverse, -m union, -a 20. Differentially expressed genes (FDR < 0.05; log_2_FC > 1) were identified using edgeR (Robinson et al., 2010) with tagwise dispersions (prior.df= 12), and genes with more than 1 CPM in at least three samples were retained for the analysis. Summary results of the inhibitor study are given in Supplemental Dataset 6.

## Accession Numbers

RNA-Seq reads used in this study were submitted to the National Center for Biotechnology Information (NCBI) Sequence Read Archive under accession numbers SRP136022 and SRP136034 (imazapyr study).

## Supplemental Data

**Supplemental Figure 1.** Distribution of gene ontology (GO) terms associated with the 1087 common DEGs.

**Supplemental Figure 2.** Differential gene expression analyses between low- and high-acylsugar-producing accessions.

**Supplemental Figure 3.** Expression levels of ten selected ‘housekeeping genes’ in different biological groups.

**Supplemental Figure 4.** Biosynthesis of branched-chain and straight-chain acyl molecules.

**Supplemental Figure 5.** Single transcript produced from the locus containing *Sopen12g004230-Sopen12g004240.*

**Supplemental Figure 6.** Correlation among expression profiles of putative defense genes.

**Supplemental Figure 7.** Lys and Arg content of selected proteins encoded by genes co-expressed with *Sopen11g004040* (FLOWERING LOCUS D; amine oxidase domain).

**Supplemental Figure 8.** Full list of enriched gene ontology (GO) terms associated with genes strongly correlated with the UDP-Glc:FA GT and the SCPL GAT.

**Supplemental Figure 9.** Top-50 most strongly connected genes for each of the three AP2-family transcription factor genes.

**Supplemental Figure 10.** Selected *O*-methyltransferase genes on chromosome 6 involved in flavonoid metabolism in *S. pennellii* and *S. lycopersicum.*

**Supplemental Table 1.** Amount of acylsugars produced by different accessions of S. *pennellii.*

**Supplemental Table 2.** Read mapping (TopHat2 alignment) results with different accessions of *S*. *pennellii.*

**Supplemental Table 3.** Sequence similarity between Cuphea KAS IV/ KAS II-like enzymes and Sopen12g004230-Sopen12g004240.

**Supplemental Table 4.** Selected genes strongly correlated with *Sopen11g004040* (FLOWERING LOCUS D; amine oxidase domain).

**Supplemental Table 5.** Selected genes in WGCNA ‘green’ module showing strong correlation with the SCPL GAT.

**Supplemental Table 6.** Selected housekeeping genes and translation-related genes showing correlations in gene expression profiles (GS) with the SCPL GAT.

**Supplemental Table 7.** Effect of imazapyr treatment on expression levels of ABC transporter genes.

**Supplemental Table 8.** Effect of imazapyr treatment on expression levels of selected genes strongly correlated with FLOWERING LOCUS D gene *Sopen11g004040* (Supplemental Table 4 genes).

**Supplemental Dataset 1.** Expression levels of all S. *pennellii* annotated genes and results of differential gene expression analysis between low- and high-acylsugar-producing accessions.

**Supplemental Dataset 2.** Spearman’s rank correlation coefficient (SRCC) among selected genes’ expression profiles in 29 biological samples (different accessions of S. *pennellii*).

**Supplemental Dataset 3.** Reciprocal best hits (RBH) between *S. pennellii* and *S. lycopersicum* annotated protein sequences.

**Supplemental Dataset 4.** Results of weighted gene correlation network analysis (WGCNA).

**Supplemental Dataset 5.** List of DEGs (between low- and high-acylsugar-producing accessions) that showed response to imazapyr at both 0.1 mM and 1 mM.

**Supplemental Dataset 6.** Summary of imazapyr treatment study.

## Author contributions

SM designed experiments, prepared samples, analyzed data, and wrote the manuscript. WJ designed experiments, prepared samples, and reviewed the manuscript. TDM designed experiments, analyzed data, and wrote the manuscript. All authors read and approved the final manuscript.

## Acknowledgements

We thank Dr. Charlie Johnson and the staff of the Texas A&M Genomics and Bioinformatics center for performing Illumina sequencing. We also thank Dr. Rodolfo Aramayo and Ricardo Perez for technical support on the BioGalaxy server. Most of the sequence analysis was performed at the Texas A&M University High Performance Research Computer, where Dr. Michael Dickens provided invaluable assistance. We sincerely appreciate Dr. Alan Pepper’s assistance with multiple aspects of this project, from initial design to critical review of the manuscript.

